# Reduced Filaggrin expression induces dysregulated intracellular signalling in atopic eczema

**DOI:** 10.1101/2024.03.11.584344

**Authors:** AJ Hughes, L Coppock, T Tachie-Menson, BR Thomas, P Dewan, I Doykov, K Mills, EA O’Toole, RFL O’Shaughnessy

## Abstract

Atopic eczema (AE) is the most common inflammatory dermatosis, affecting up to 20% of children. Loss of function mutations in the Filaggrin (*FLG*) gene are the most strongly implicated genetic risk factor for AE, but little is known about the signalling pathways altered in response to loss of *FLG.* To explore the downstream effects of loss of FLG on the cellular environment, we combined RNAseq analysis of siRNA knockdown of normal human keratinocytes and analysis of tape strip (TS) samples from AE patients who were clinically phenotyped and genotyped for *FLG* mutation status.

RNA-seq analysis revealed an increase in BMP signalling following FLG KD, which we validated *in vivo* using TS samples and biopsies. Recombinant BMP2 or BMP6 increased FLG and suprabasal keratin expression in vitro. Phosphoproteomic analysis identified 237 significantly differentially phosphorylated proteins following FLG KD. Kinase enrichment analysis identified downregulation of ERK1/2 and AKT1 signalling which was confirmed in AE biopsies. cFOS was downregulated following FLG KD and correlated with FLG expression in repository datasets. cFOS was downregulated following BMP6 treatment, implying that cFOS may be an important link between FLG and BMP signalling. Finally, proteomic analysis of TS samples identified altered desmosomal expression and phosphorylation following either loss of FLG or increased BMP signalling. Therefore, we have identified a SMAD1/Filaggrin/AKT axis as a potential therapeutic avenue in AE.

## Introduction

Atopic eczema (AE) is the most common inflammatory dermatosis and affects between 10-20% of children (Abuabara et al., 2019; Odhiambo et al., 2009; Paternoster et al., 2018), with persistence into adulthood in 50% of cases (Abuabara et al., 2019). AE is a complex trait disease and the hereditary component is estimated at 75% (Elmose & Thomsen, 2015). Variants in the Filaggrin gene (*FLG*) are the most strongly implicated genetic risk factor for AE (Bin & Leung, 2016), but account for only 20% of AE patients in the white European population (Brown et al., 2009; Henderson et al., 2008; Palmer et al., 2006). In other ethnic populations *FLG* mutations affect up to 50% of individuals (Pigors et al., 2018).

The Filaggrin protein (FLG) undergoes complex processing. It is synthesised as the inactive macroprotein profilaggrin in the stratum granulosum, then processed by a cascade of proteases to the active 37 kDa FLG monomer (de Veer et al., 2014). FLG later undergoes deimination and further proteolytic processing to natural moisturising factors (NMF) (Cau et al., 2017). The NMF bind water to confer hydration to the skin, protect the skin from UV light and have an anti-inflammatory role (Rieko & Motonobu, 2016; Scott & Harding, 1986; Simonsen et al., 2017). The prevailing opinion is that the positively charged FLG monomers bind to negatively charged suprabasal keratins, collapsing the keratin network and binding these proteins together to form a part of the stratum corneum (Drislane & Irvine, 2020; Kypriotou et al., 2012). However, there is little data to support this *in vivo* and *in vitro* this only occurs when there is forced expression of FLG (Dale et al., 1997). During terminal differentiation, FLG translocates towards the nucleus (Gutowska-Owsiak et al., 2018) and part of the FLG protein, the N-terminus, enters the nucleus (Pearton et al., 2002). FLG is therefore thought to play a role in triggering nuclear degradation and terminal differentiation. Knock down (KD) of the *FLG* gene has been shown to affect signalling pathways within primary keratinocytes (S. Wang et al., 2018). Attempts to therapeutically directly restore or replace FLG so far have been unsuccessful, for example using readthrough drugs (Irvine, 2014), therefore there is increased interest in understanding signalling both upstream and downstream of FLG expression. In this paper, we demonstrate that loss of FLG has a profound effect on the intracellular signalling environment, increasing BMP signalling and reducing activity of AKT and MAP kinases, providing new AE therapy targets.

## Materials and Methods

### Cell culture

P0 NHEKs isolated from neonatal foreskin (Thermo Fisher, USA) were cultured on a feeder layer of mitomycin treated J2 Clone of Swiss 3T3 Cells (J2-3T3s) in FAD media (3 parts Dulbecco’s Modified Eagle medium (Thermofisher, USA), 1 part Ham’s F12 media mixture (Thermofisher, USA), 10% FBS (Thermofisher, USA), 200mM L-glutamine (Lonza, Switzerland), 1% penicillin-streptomycin (Thermofisher, USA), 5 μg/mL Insulin (Sigma-Aldrich, USA), 0.18 mM Adenine (Thermofisher, USA), 0.5 μg/mL Hydrocortisone (Thermofisher, USA), 0.1 nM Cholera Toxin (Sigma-Aldrich, USA), 10 ng/ml EGF (Peprotech, USA)). Y-27632 (Cell Guidance Systems, Cambridge, UK), a Rho Kinase Inhibitor was added to the media at a final concentration of 10 μM to prevent terminal differentiation during passage and removed at least 10 days before analyses were performed. NHEKs were incubated at 37°C with 5% CO_2_ in a humidified atmosphere. Media were changed every 2-3 days, NHEKs were split when colonies were subconfluent approximately every 7-10 days. Rat epidermal keratinocytes (REKs) were grown as previously described (Rogerson et al., 2021).

### Western blot and antibodies

NHEKs and REKs were lysed using a lysis buffer as previously described (Tagoe et al., 2023). For NHEKs, J2-3T3 cells were removed using a single versene wash prior to lysis. Western blots were run and immunoblotting performed as previously described (Tagoe et al., 2023). The following antibodies were used: Anti-Filaggrin AE21(Santa Cruz Biotechnology Inc, USA) 1:750, Anti-Filaggrin AKH1(Santa Cruz Biotechnology Inc, USA) 1:200, Anti-Keratin 1 905601(Biolegend, USA) 1:1000, Anti-Keratin 2 bs-1005R (Bioss, USA) 1:500, Anti-Keratin 10 905404 (Biolegend, USA) 1:1000, Anti-Phospho-SMAD1(Ser463/465)/ SMAD5(Ser463/465)/ SMAD9(Ser465/467) (D5B10) 13820 (Cell Signalling Technologies, USA) 1:1000, Anti-SMAD 1 10429-1-AP (Proteintech, USA) 1:500, Anti-SMAD 1 66559-1-Ig (Proteintech, USA) 1:4000, Anti-Loricrin ab85679 (Abcam, UK) 1:1000, Anti-BMP2 bs-1012 (Bioss, USA) 1:300, Anti-ID1 18475-1-AP (Proteintech, USA), 1:500, Alpha tubulin ab7291 (Abcam, UK) 1:5000, Anti-cFos 2250S (Cell Signalling Technologies, USA) 1:1000, Anti-Phospho-AKT (Ser473) 9271 (Cell Signalling Technologies, USA) 1:1000, Anti-AKT (pan) (C67E7) 4691 (Cell Signalling Technologies, USA) 1:1000, Anti-Phospho-p44/42 MAPK (Erk1/2) (Thr202/Tyr204) (E10) (Cell Signalling Technologies, USA) 1:1000, Anti-p44/42 MAPK (Erk1/2) 9102 (Cell Signalling Technologies, USA) 1:1000, Anti-Cathepsin H (F7) sc-398527(Santa Cruz Biotechnology Inc, USA) 1:500, Anti-Phospho-cFos (D82C12)(Ser32) 5348 (Cell Signalling Technologies, USA) 1:1000, Anti-SPRR3 11742-1-AP (Proteintech, USA), 1:750, Anti-Repetin bs-9181R (Bioss, USA) 1:500, Anti-GAPDH MAB374 MilliporeSigma,USA), 1:3000, Anti-Filaggrin 2 (Cloud-Clone, USA), 1:750, Anti-S100A14 10489-1-AP (Proteintech, USA), 1:500, Anti-BMP7 12221-1-AP (Proteintech, USA) 1:500, Anti-p38 MAPK (D13E1)8690 (Cell Signalling Technologies, USA) 1:1000, Anti-Phospho-p38 MAPK (Thr180/Tyr182) (D3F9) 4511 (Cell Signalling Technologies, USA) 1:1000.

### FLG shRNA knockdown (KD) of NHEKs

Tetracycline inducible SMARTvector 2.0 microRNA expression scaffold with CMV promoter, Turbo reporter, and puromycin selection cassette (Horizon Discovery, UK). Four different constructs were used: V3SH7669-225732812, V3SH7669-226710173, V3SH7669-228900284 and a non-targeting mCMV-TurboRFP Control (VSC6571). 2×10^6^ puromycin-resistant mitomycin treated J2-3T3 cells were seeded into a 10 cm dish on Day 0. NHEKs were seeded on Day 1 at 50,000 per dish in FAD media. On Day 2 the media were changed to FAD with polybrene 2.5 µg/ml. Lentivirus was added to an MOI of 0.5. NHEKs were grown in FAD with puromycin 1 µg/ml for 7 days to select the virally transduced cells. KD was confirmed using qPCR and western blotting. 12,500 *FLG* shRNA KD NHEKs were seeded onto mitomycin treated J2-3T3s in 6-well plates and grown for 2 weeks. 1 µg/ml of doxycycline was added to the media from day 4. After 2 weeks the cells were prepared either for western blot or RNA extraction as detailed above.

### *FLG* siRNA KD of NHEKs

Custom made siRNA (Invitrogen, UK) sequence-specific to FLG was transfected into NHEKs, sequence: ACAGAAAGCACAGUCAUCAUGAUAA, along with a non-targeting siRNA control NHEKs were plated at a density of 1.5×10^5^ per well and incubated in Epilife media (Cascade Biologics, UK) After 24 hours, NHEKs were transfected with FLG siRNA and the non-targeting control using HiPerFect (Qiagen,UK). For transfection, siRNA duplexes (5-25nM) and 15 μl of HiPerFect transfection reagent were diluted with 95 μl Epilife media, transfection as per manufacturers instuctions. The following day transfection media were replaced with Epilife. 12 hours post-transfection, the cells were shifted to high calcium (1.3 mM) and incubated for 24 hours. Cells incubated in low calcium media for 24 hours were kept as a control. RNA was extracted 3 days post-calcium switch.

### *Flg* siRNA KD in REKs

FlexiTube siRNA Premix siRNA (Qiagen, Germany): SI01513407|S5, SI01513414|S5, SI01513421|S5 and a scrambled control (SI03650325|S5) were used. REKs were plated at 0.4×10^5^ cells in a 12 well plate. The following day, 25 µL siRNA was added. 24 hours later the media were changed. 48 hours later, once at post confluency, the cells were either lysed for protein extraction or RNA extraction. Alternatively, REKs were trypsinized and plated onto collagen coated cover slips, then fixed and stained for immunocytochemistry. *Flg* KD was confirmed using western blotting and qPCR.

### Recombinant BMP, FST and DMH1 Experiments

40,000-50,000 REKs were plated in a 12-well plate with DMEM+ media with 10% FCS. 6 hours later the media were changed to DMEM+ media without serum. The following day recombinant BMP2 (rBMP2)(Biolegend, USA), recombinant BMP6 (rBMP6) (Biolegend, USA), recombinant Follistatin (rFST) (Biolegend, USA) or vehicle control were added to the desired concentration whilst still preconfluent (approximately 24 hours before confluency). Concentrations of 25, 125 or 200 ng/mL of rBMP2 and rBMP6 were used. rFST concentrations were 50, 250 and 1000 ng/mL. DMH1 (Sigma-Aldrich, USA) was used at a dose of either 10 µM or 20 µM. 24 hours after recombinant protein treatment the media was changed to DMEM+ with serum. The cells were cultured for 2 days and lysed.

### RNA Extraction

RNA was extracted using the RNEasy Mini Plus kit (Qiagen, Germany). For NHEKs the J2s were removed with a single 5-minute wash with versene prior to lysis. RNA was quantified using spectrophotometry (Nanodrop, ND-100 Spectrophotometer, USA). For qPCR, cDNA was first synthesised using the Lunascript RT Supermix (New England Biolabs, USA). qPCR was performed on the synthesised cDNA using the Luna Universal qPCR Master Mix kit (New England Biolabs, USA). 4 μL of cDNA was added to 1 μL of pre-purchased primers (QuantiTect primers (Qiagen, Germany)) and qPCR was carried using the StepOnePlus machine (Thermo Fisher Scientific. USA).

### RNAseq Analysis

RNA was extracted from *FLG* siRNA KD post confluent cells as detailed above, then sent to GSK (RD Molecular Discovery Research,GlaxoSmithKline, Collegeville, PA, USA) for sequencing. All samples had an RNA Integrity Number of >8. RNA was converted to a library of cDNA fragments and adaptors attached to one or both ends. Following amplification each molecule was pair-end sequenced in a high-throughput manner using the Illumina platform The reads were not strand specific. KRAKEN (http://www.ebi.ac.uk/research/enright/software/kraken) pipeline, was used to process the RNA-Seq data. First, the 3’ adaptor sequences were trimmed. Then, identical reads were made to a single entry while keeping the depth values. A file with unique reads and their corresponding counts was generated for each sample. The final processed unique paired-end reads were aligned to the reference human genome (GRCh37/hg19) with Tophat v2.0.4. For transcript quantification, the number of reads mapped to transcripts (Homo_sapiens.GRCh37.74.gtf from Ensembl) were obtained from Tophat2 aligned sam files using the HTSeq-count command from the HTSeq library (http://www.huber.embl.de/users/anders/HTSeq/doc/). HTSeq was used to assemble individual transcripts from the reads. Read count, normalizion and differential expression was calculated using the Bioconductor DESeq2 package (http://www.bioconductor.org/packages/release/bioc/html/DESeq2.html). Differential gene expression analysis was based on a negative binomial distribution.

### Immunofluoresence

For immunocytochemistry, REKs were grown to confluency, trypsinised and replated onto coverslips. Fixation, blocking, staining and microscopy were perfomed as previously described (Rogerson et al., 2021). The following antibodies were used: Anti-Phospho-SMAD1(Ser463/465)/ SMAD5(Ser463/465)/ SMAD9(Ser465/467) (D5B10) 13820 (Cell Signalling Technologies, USA) 1:1000.

### Phosphoproteomic Analysis

FLG shRNA KD NHEKs were grown to post confluency for a total of 14 days. Cells were placed on ice and washed 3 times with 2 mL ice cold PBS containing phosphatase inhibitors (1mM sodium fluoride and 1mM sodium orthovanadate), then lysed using a buffer of 8M urea in 20mM HEPES, pH 8.0 and phosphatase inhibitors (1mM sodium orthovanadate, 1mM sodium fluoride, 1mM beta-glycerol phosphate and 2.5mM disodium pyrophosphate. The lysate was placed in a lo-bind Eppendorf 1.5 mL tube then sonicated on ice, using a Bioruptor sonicator (Diagenode, Belgium). 10x 30 second cycles of sonication were used on a high setting, with a 30 second interval between cycles. The lysates were centrifuged for 10 minutes at 13,000 RPM at 4°C. The supernatant was quantified using the DC protein assay (Bio-Rad Laboratories Inc, USA). 110 µg of protein was placed into a low-bind Eppendorf 1.5ml tube and the volume was normalised to 200 µl using the lysis buffer.

Proteins were digested into peptides using trypsin as previously described (Alcolea et al., 2012; Montoya et al., 2011). Phosphopeptides were desalted and enriched using the AssayMAP Bravo (Agilent Technologies) platform. For desalting, protocol peptide clean-up v3.0 was used. Reverse phase-S cartridges (Agilent, 5 μL bed volume) were primed with 250 μL 99.9% acetonitrile (ACN) with 0.1%TFA and equilibrated with 250 0.1% TFA at a flow rate of 10 μL/min. The samples were loaded at 20 μL/min, followed by an internal cartridge wash with 0.1% TFA at a flow rate of 10 μL/min. Peptides were then eluted with 105 μL of 1M glycolic acid with 50% ACN, 5% TFA and this is the same buffer for subsequent phosphopeptide enrichment. Following the Phospho Enrichment v 2.1 protocol, phosphopeptides were enriched using 5ul Assay MAP TiO2 cartridges on the Assay MAP Bravo platform. The cartridges were primed with 100ul of 5% ammonia solution with 15% ACN at a flow rate of 300 μL/min and equilibrated with 50 μL loading buffer (1M glycolic acid with 80% ACN, 5% TFA) at 10 μL/min. Samples eluted from the desalting were loaded onto the cartridge at 3 μL/min. The cartridges were washed with 50 μL loading buffer and the phosphorylated peptides were eluted with 25 μL 5% ammonia solution with 15% ACN directly into 25 μL 10% formic acid. Phosphopeptides were lyophilized in a vacuum concentrator and stored at -80°C.

Dried phosphopeptides were dissolved in 0.1% TFA and analysed by nanoflow ultimate 3000 RSL nano instrument was coupled on-line to a Q Exactive plus mass spectrometer (Thermo Fisher Scientific). Gradient elution was from 3% to 28% solvent B in 90 min at a flow rate 250 nL/min with solvent A being used to balance the mobile phase (buffer A was 0.1% formic acid in water and B was 0.1% formic acid in acetonitrile) . The spray voltage was 1.95 kV and the capillary temperature was set to 255 °C. The Q-Exactive plus was operated in data dependent mode with one survey MS scan followed by 15 MS/MS scans. The full scans were acquired in the mass analyser at 375-1500m/z with the resolution of 70 000, and the MS/MS scans were obtained with a resolution of 17 500.

MS raw files were converted into Mascot Generic Format using Mascot Distiller (version 2.8.1) and searched against the SwissProt database (SwissProt_2021_02) restricted to human entries using the Mascot search daemon (version 2.8.0). Allowed mass windows were 10 ppm and 25 mmu for parent and fragment mass to charge values, respectively. Variable modifications included in searches were oxidation of methionine, pyro-glu (N-term) and phosphorylation of serine, threonine and tyrosine. Phosphopeptide quantification was performed using in-house software Pescal as described before (Alcolea et al., 2012). The resulting quantitative data was parsed into excel files for further normalisation and statistical analysis.

The differential expression of phosphopeptides was filtered to include only results where there was a log2fold change of >/ 0.58 or /<-0.58 and p value of <0.05 relative to the control. Proteins were identified from the differentially expressed phosphopeptides and converted to their corresponding gene symbols. Gene symbols were uploaded into the KEA3 online platform (available from https://maayanlab.cloud/kea3), to predict the over-representation of upstream kinases from an uploaded dataset by integrating 11 kinase-substrate libraries covering 520 unique protein kinases, protein-protein interaction databases as well as datasets that utilise co-occurrence, and transcript co-expression data (Kuleshov et al., 2021).

### Patient Samples and Clinical Data

Bangladeshi children and young adults age 0-30 with AE were recruited as part of a wider study at the Royal London Hospital and underwent clinical phenotyping and *FLG* genotyping as previously described (Thomas et al., 2023). This study was approved by the local ethics committee: Hampstead Regional Ethics Committee reference 18/LO/0018; Patients and/or parents gave written informed consent before participation in the study. Patients aged 14-30 were invited to return for a 3mm punch biopsy. Biopsies were taken from non-lesional skin on the volar forearm under local anaesthetic. Retrospective patient notes review was used to identify serum vitamin D3 levels and were included only if these had been taken previously.

### Tissue Section Staining

Immunofluorescence of tissue samples was performed as previously described (A. S. Naeem et al., 2015; Aishath S. Naeem et al., 2017; Rogerson et al., 2021). The following antibodies were used: Anti-Phospho-AKT (Ser473) 9271 (Cell Signalling Technologies, USA) 1:25, Anti-Phospho-p44/42 MAPK (Erk1/2) (Thr202/Tyr204) (E10) (Cell Signalling Technologies, USA) 1:50, Anti-cFos 2250S (Cell Signalling Technologies, USA) 1:50, Anti-BMP2 bs-1012 (Bioss, USA) 1:50, Anti-ID1 18475-1-AP (Proteintech, USA) 1:100, Anti-Filaggrin (Biolegend, USA) 1:50.

### Tape Stripping Protocol

22.0mm D-squame® D100 brand tape strips were used with a D500 D-squame® pressure instrument (Clinical & Derm, USA). TS were applied to non-lesional skin on the volar forearm. A standardised pressure (225g/cm^2^) was applied using the pressure instrument for 10 seconds before removing the strip with forceps. Marker pen was used to draw around the first TS and the first TS was discarded to remove any superficial contaminants. TS 2-20 were collected from patients and pooled into superficial (numbers 2-10) and deep TS (numbers 11-20).

For western blot analysis, superficial TS were cut into quarters. The set of quarters were split between four 1.5 mL centrifuge tubes (9 per tube) and placed on ice. 600 µl of lysis buffer containing 1X PBS with 0.2% SDS and 1mM DTT was added to each tube. Tubes were sonicated on ice for 15 minutes on a high setting with a 30 second interval every 30 seconds. Ice was changed every 10 minutes. The samples were heated for 30 minutes at 95°C with the TS left in the tube, then centrifuged at 13000 RPM for 10 minutes. Protein was quantified using the DC Protein Assay (Bio-Rad Laboratories Inc, USA). Samples were divided into aliquots and stored at -20°C. On thawing x6 Laemmli loading buffer was added (at a ratio of 1:5) and the samples were heated to 95°C for 1 minute.

Equal protein was loaded (8µg for less abundant proteins, 5µg for more abundant proteins) onto a precast protein gel. A non-AE control sample was loaded to each gel from the same individual to allow for comparison between samples. Again western blots were performed as previously described (Tagoe et al., 2023)

### Mass Spectrometry Proteomics of TS Samples

TS 2-20 were separated and cut into quarters. Half of TS 2-10 were reserved for protein extraction for western blotting. The other half of TS 2-10, together with TS 11-20 were placed into x3 1.5 mL centrifuge tubes. 600 µL of a lysis buffer containing 100 mM Tris HCl, pH 7.2, 8M urea and 1mm DTT was added to each tube. Samples were sonicated and centrifuged as detailed above. Lysates from each set of TS were pooled together, frozen on dry ice and kept at -80°C until MS proteomic analysis. Protein fractions were purified by acetone precipitation and prepared for mass spectrometry analyses according to (Toomey et al 2022, n=14). Label free quantitative proteomic analyses was carried out on nano LC-Cyclic QTOF (Waters, UK) as previously described (Hällqvist et al., 2023). Data acquired were analysed using Progenesis LC–MS (Nonlinear Dynamics Limited, Newcastle, UK) raw data was processed as described previously but with the Uniprot database (June 2023) [24]. Briefly peptides were searched as described above except with 1% false discovery rate. Protein data for identifications with a confidence score > 20 and more than one unique peptide were exported for further analysis.

### Bioinformatics

As a label-free approach was taken, all genes or proteins that met the threshold set (> twofold change in one region, and also changing in four other regions) were subject to gene ontology analysis using Ingenuity Pathway Analysis software (Qiagen) was used to perform in depth canonical pathway analysis and determine biological functions altered in the datasets. Multivariate analysis on the second data analysis on the Synapt G2 MS was performed using SIMCA v15 (Umetrics, Sweden).

### Microscopy

Epi-fluorescence and brightfield images were taken on the Leica DM5000B Epi-Fluorescence microscope. Confocal images were taken on the Zeiss 710 Laser Scanning Confocal microscope.

### Repository Bioinformatic Analysis

Transcriptomic repository datasets for patients with atopic dermatitis were analysed by accessing the website https://www.ncbi.nlm.nih.gov/. The search term ‘atopic dermatitis’ was inputted into the category ‘GEO DataSets’ and filtered to include only ‘Expression Profile by Array’ under the Study Type category. Only microarray studies were included as these have already been mapped to the reference genome and normalized, allowing a larger number of datasets to be analysed. Samples were included only if there was a minimum of 10 patient samples. Both lesional and non-lesional samples were included. Samples were not included if there was an intervention applied (eg a drug treatment). Untreated samples were included if the sample size was large enough. If there was a significant correlation between FLG and Th inflammatory components (IL4, IL13, or STAT1) within the dataset only non-lesional samples were included. If there were < 10 non-lesional samples the study was excluded. This is because Th2 cytokines are known to reduce FLG expression and may have confounded the analysis (Hönzke et al., 2016; Howell et al., 2009). Different microarray datasets are not directly comparable due to different sequencing platforms, study design and experimental variables (Choi et al., 2003). For each dataset, associations were determined for FLG expression relative to other EDC, Keratin and BMP genes and plotted as scatter plots. Correlation coefficients were calculated by using simple linear regression, calculated using Prism 9. A heatmap was used to compare studies. The following studies were included from the platform: GDS4444, GDS4491, GSE1037361, GSE58558, GSE111055, GSE120721.

### Statistical Analysis

Biological triplicates were used for the RNAseq, phosphoproteomic analysis, immunocytochemistry staining, and for western blots. Technical triplicates were used for qPCR experiments. Simple linear regression was performed for correlation analysis. For qPCR, immunostaining and western blots from patient samples, the one-way ANOVA with Dunnet’s post-hoc testing was performed. Prism v9 software (GraphPad software Inc, USA) was used for statistical analysis.

Volcano plots were made using the function on the website www.usegalaxy.org

Differentially expressed proteins were also uploaded to STRING for analysis (accessed from https://string-db.org) and visualised as a physical subnetwork. The settings used were physical subnetwork, medium confidence (0.400) with disconnected nodes removed.

Differentially expressed gene IDs were exported to the Enrichr web application (available from https://maayanlab.cloud/Enrichr) for transcription factor enrichment (E. Y. Chen et al., 2013).

## Results

### RNAseq analysis of Filaggrin knockdown keratinocytes revealed widespread downregulation in epidermal terminal differentiation genes

We performed RNA-seq analysis to globally assess changes to the transcriptome and establish differential pathway expression following FLG siRNA knockdown (KD). Over 6000 differentially expressed genes were identified with a marked reduction in several genes related to the cornified envelope and terminal differentiation (Figure 1a and b). We confirmed downregulation of terminal differentiation proteins following FLG KD by western blotting and qPCR (Figure 1c and d). We also confirmed this in Flg shRNA knockdown of rat epidermal keratinocytes (REKs) (Supplementary Figure S1a-c). Gene expression omnibus AE microarray datasets showed a consistent positive correlation between FLG and terminal differentiation genes in all but 1 dataset (Figure 1e). Tape strip (TS) samples were used to study components of the stratum corneum in Bangladeshi patients from the East end of London with AE (Figure 1f). The Bangladeshi AE endotype has a large number of FLG LoF mutations and enrichment in variation of FLG mutations (Pigors et al., 2018). The positive correlations between FLG and many terminal differentiation proteins demonstrated *in vitro* and *in silico* were not replicated *in vivo* (Supplementary Figure S2a) but there a positive correlation was identified between KRT1/KRT2 and FLG (Figure 1f). Overall, this suggests that *in vivo*, other factors alter the expression of terminal differentiation proteins.

**Figure 1:**
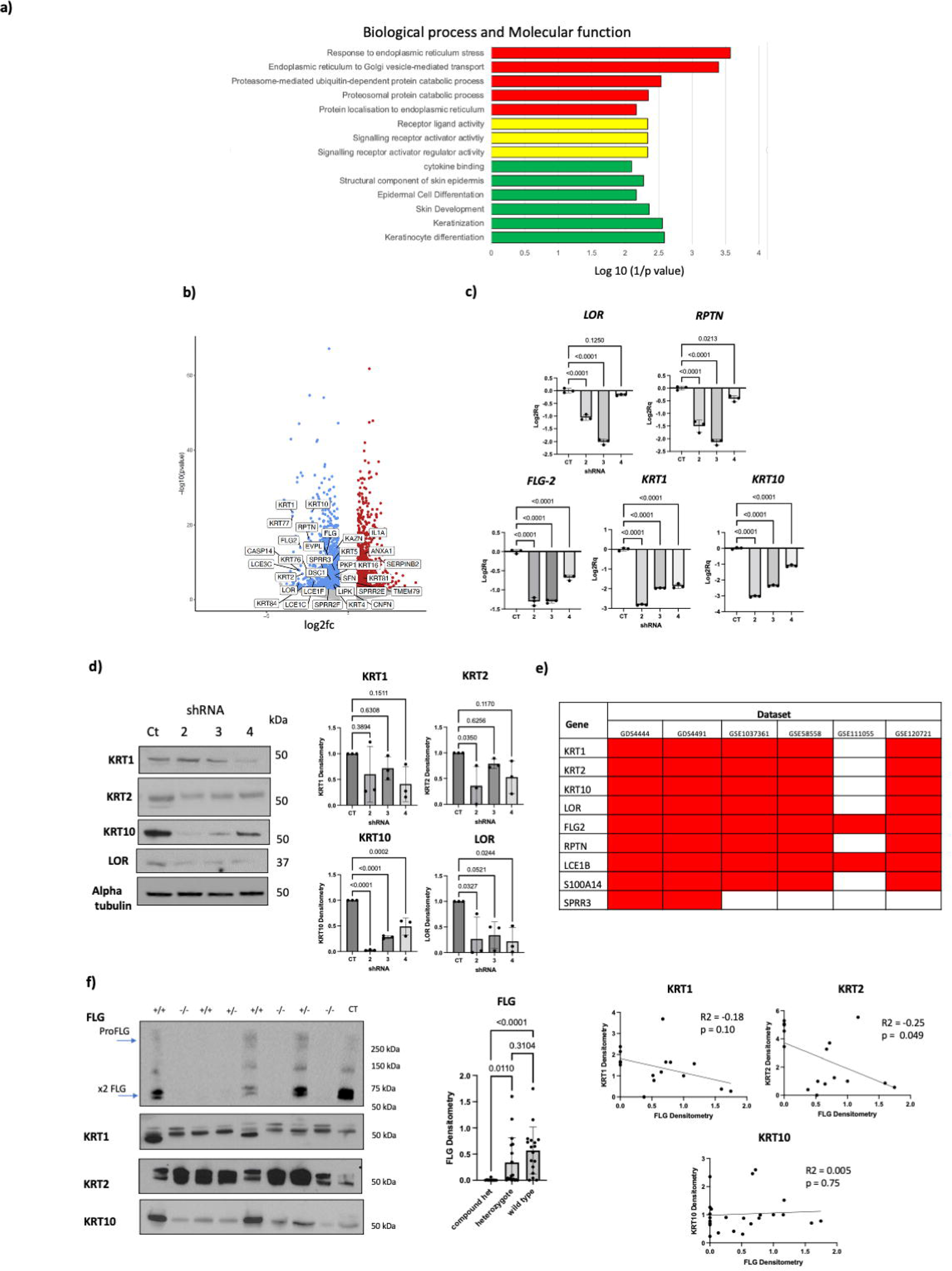
FLG Knock Down Causes Loss of Terminal Differentiation Proteins and Dysregulation of Intracellular Signalling Pathways. a) Gene expression omnibus analysis of the FLG siRNA KD RNA-seq dataset showing the top 10 significant terms for cellular components and all the significant terms for molecular function. b) Volcano plot showing differentially expressed terminal differentiation genes using RNA-seq following FLG siRNA KD. c) qPCR following FLG shRNA KD in NHEKs expressed as Log2Rq relative to control. d) Western blots following FLG shRNA KD in NHEKs (representative of 3 replicates). e) Correlation analysis of FLG with EDC and terminal differentiation genes from AE patient microarray repository datasets. Red = significant positive correlation. Green = significant negative correlation. f) Western blots using AE patient tape strip lysates. +/+ = wild type, +/-= heterozygote, -/-= compound heterozygote. g) scatter plots showing the relationship between FLG and suprabasal KRTs.

We analysed the clinical data to try to determine which exogenous factors may account for this difference. We found a positive correlation between FLG expression and serum vitamin D3 and a negative correlation between KRT2 and serum vitamin D3 (Supplementary Figure S2b). We hypothesise that vitamin D3 has a confounding effect on the relationship between FLG and suprabasal keratin expression *in vivo*. Vitamin D3 has previously been shown to reduce keratinocyte proliferation and induce differentiation in vitro (Bikle et al., 1993). This may be more relevant to Bangladeshi AE, as people of colour in northern latitudes have a high risk of vitamin D3 deficiency (Darling et al., 2013; Smith et al., 2021).

### BMP signalling is increased in Filaggrin knockdown keratinocytes and in eczema gene expression datasets. Increased BMP signalling induced increased filaggrin expression and processing

We reviewed the FLG KD RNA-seq dataset to identify other downstream factors that might be applicable to the population we were studying. We were interested that loss of FLG caused changes to signalling receptor binding (Figure 1a). Filaggrin is thought to be a structural protein and has not previously been associated with changes to signalling pathways. We were particularly interested in the changes to ligands involved in bone morphogenetic protein (BMP) signalling (Figure 2a) following FLG KD, particularly the increase in BMP6 with a corresponding downregulation of its inhibitor noggin (NOG) and the ligand follistatin (FST) (Figure 2a). BMP ligands have previously been shown to induce the expression of terminal differentiation genes (Gosselet et al., 2007; Kim et al., 2018; McDonnell et al., 2001; Phillips et al., 2013). As BMP6 and FST are both secreted ligands we hypothesised that these ligands are secreted as a direct response to an impairment in terminal differentiation, induced by a loss of FLG.

**Figure 2:**
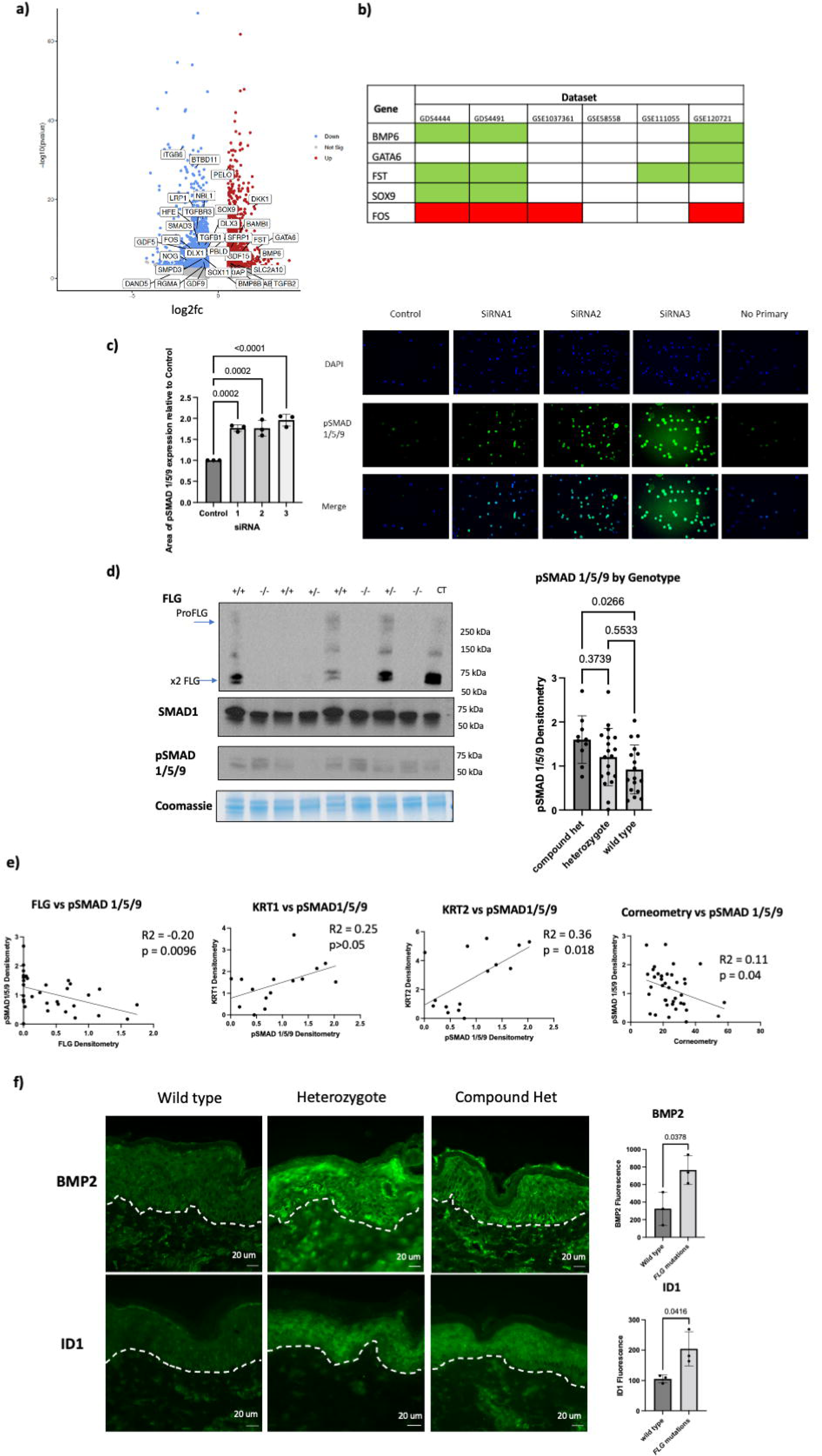
Loss of FLG Is Associated with an Increase in BMP Signalling. a) Volcano plot showing differentially expressed BMP related genes using RNA-seq following FLG siRNA KD b) Correlation of FLG with BMP related genes from AE patient microarray repository datasets. Red = significant positive correlation. Green = significant negative correlation. The top 5 BMP related genes were selected from the FLG KD RNA-seq dataset in Figure 2a. c) Cover slip staining for pSMAD 1/5/9 following Flg siRNA knock down in REKs d) Western blots using AE patient tape strip lysates. +/+ = wild type, +/-= heterozygote, -/-= compound heterozygote. d) Scatter plot correlating pSMAD 1/5/9 expression from AE patients’ tape strip lysates with FLG, KRT1, KRT2 and clinical corneometry measurements. f) Tissue sections from AE patients with staining for BMP2 and ID1.

BMP6 stimulates the canonical BMP signalling pathway through receptor binding and intracellular phosphorylation of SMAD 1/5/9 (also known as SMAD 1/5/8). It also activates the Mitogen Activated Protein Kinase (MAPK) and c-Jun-N-terminal Kinase (JNK) signalling pathways through the non-canonical BMP signalling pathway (Botchkarev, 2003; Botchkarev & Sharov, 2004; Singh et al., 2012). FST inhibits the Transforming Growth Factor-Beta (TGF-Beta) signalling pathway by binding to Activin A (Moura et al., 2013). phospho-SMAD 1/5/9 (pSMAD1/5/9), the activated ligand of SMAD1/5/9, translocates into the nucleus where it binds to promoter sites to regulate cell growth, differentiation and apoptosis (Botchkarev & Sharov, 2004; Morikawa et al., 2011). We saw an increase in pSMAD1/5/9 in Flg siRNA KD REKs (Figure 2c)

BMP6 and FST negatively correlated with FLG using repository dataset analysis (Figure 2b), but only in a subset of the studies analysed. This suggests that the association between BMP signalling and FLG expression may not be relevant in all AE populations (Czarnowicki et al., 2019), Western blots performed on tape strip (TS) lysates from AE patients showed an increase in pSMAD1/5/9 in compound heterozygous patients (with 2 *FLG* mutations) relative to wild type patients in the Bangladeshi AE cohort (Figure 2d). Overall FLG expression negatively correlated with pSMAD1/5/9 expression, suggesting patients with low FLG had increased BMP signalling (Figure 2e). Epidermal BMP2 and ID1, a downstream target of BMP signalling, were increased in patients with *FLG* mutations (Figure 2f). pSMAD1/5/9 levels also negatively correlated with corneometry, a clinical measure of skin hydration (Figure 2e). As pSMAD1/5/9 also positively correlated with suprabasal keratin expression (Figure 2e), we hypothesised that pSMAD1/5/9 induces the expression of suprabasal keratins as a response to barrier impairment.

We went on to mechanistically explore the role of BMP signalling on keratinocytes *in vitro*. Treatment with either recombinant BMP6 (rBMP6), recombinant BMP2 (rBMP2) or recombinant FST (rFST) increased FLG and KRT1 expression (Figure 3a), while treatment with the BMP receptor antagonist dorsomorphin homolog 1 (DMH1) reduced their expression (Figure 3b). These data suggest that increased BMP signalling is a protective response that induces the expression of terminal differentiation proteins to restore the skin barrier.

**Figure 3:**
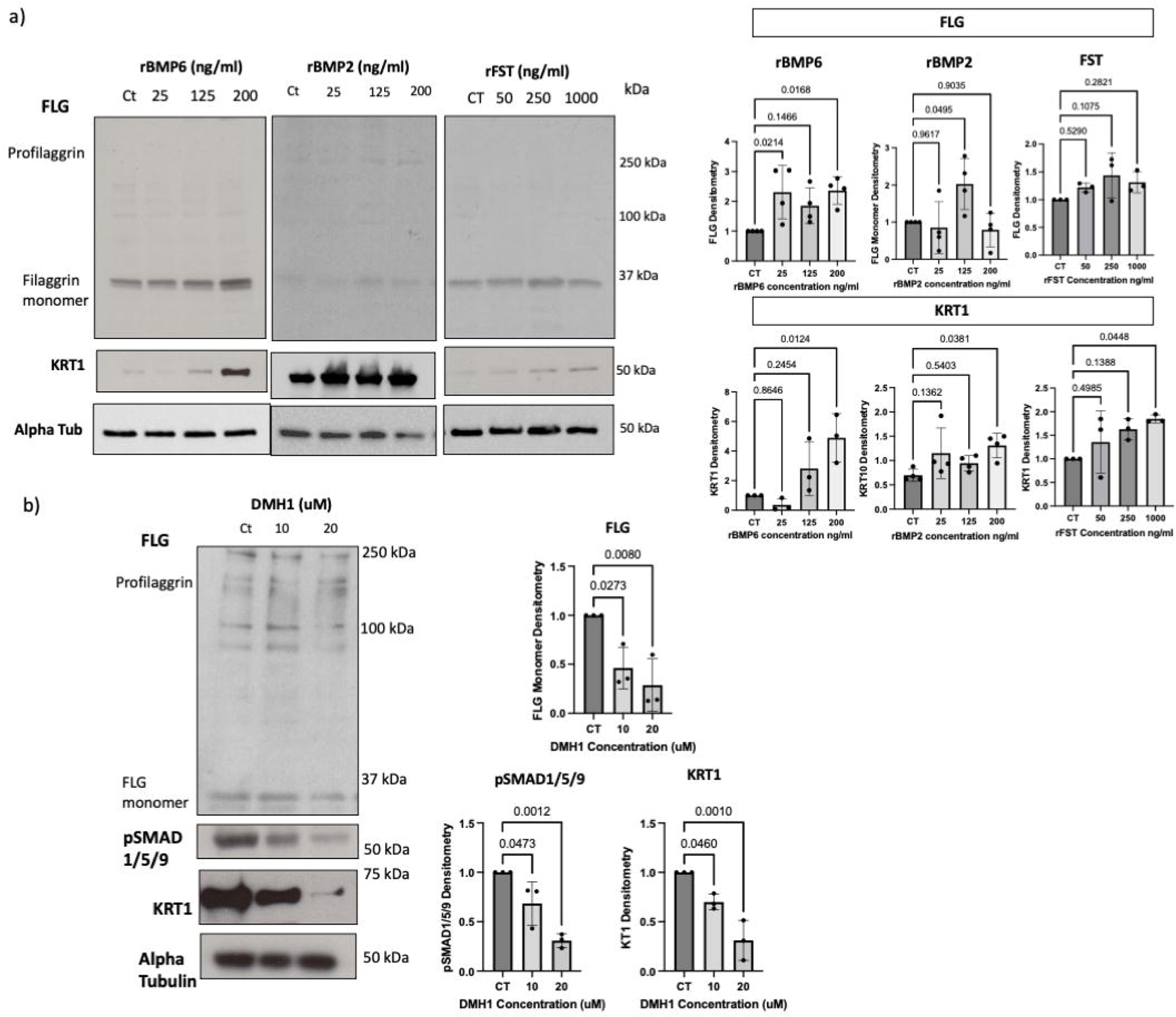
BMP2, BMP6 and FST Increase Terminal Differentiation Protein Expression in Keratinocytes. a) Western blots showing FLG and KRT1 expression following treatment of REKs with recombinant BMP6, BMP2 or FST. b) Western blots showing FLG expression in REKs following treatment with DMH1, a selective BMP-receptor antagonist.

### Filaggrin knockdown downregulated AKT and ERK kinase activity

We were interested to comprehensively explore changes to intracellular signalling pathways. Phosphoproteomic analysis identified 428 significantly differentially phosphorylated proteins following FLG KD. STRING analysis of differentially phosphorylated proteins revealed a cluster involving keratin proteins (Supplementary Figure S4a), suggesting that loss of FLG causes a reduction in keratin phosphorylation as well as expression. Actin cytoskeleton regulators were also differentially phosphorylated (Supplementary Figure S4a). This was of interest because FLG keratohyalin granules use the actin scaffold to translocate towards the nucleus prior to the onset of terminal differentiation (Gutowska-Owsiak et al., 2018). Interestingly, the FLG protein itself was differentially phosphorylated in this dataset (Supplementary Figure S4a), suggesting that FLG controls its own phosphorylation. Differential phosphorylation occurred at a linker site between FLG monomers on the profilaggrin molecule (Figure 4d).

**Figure 4:**
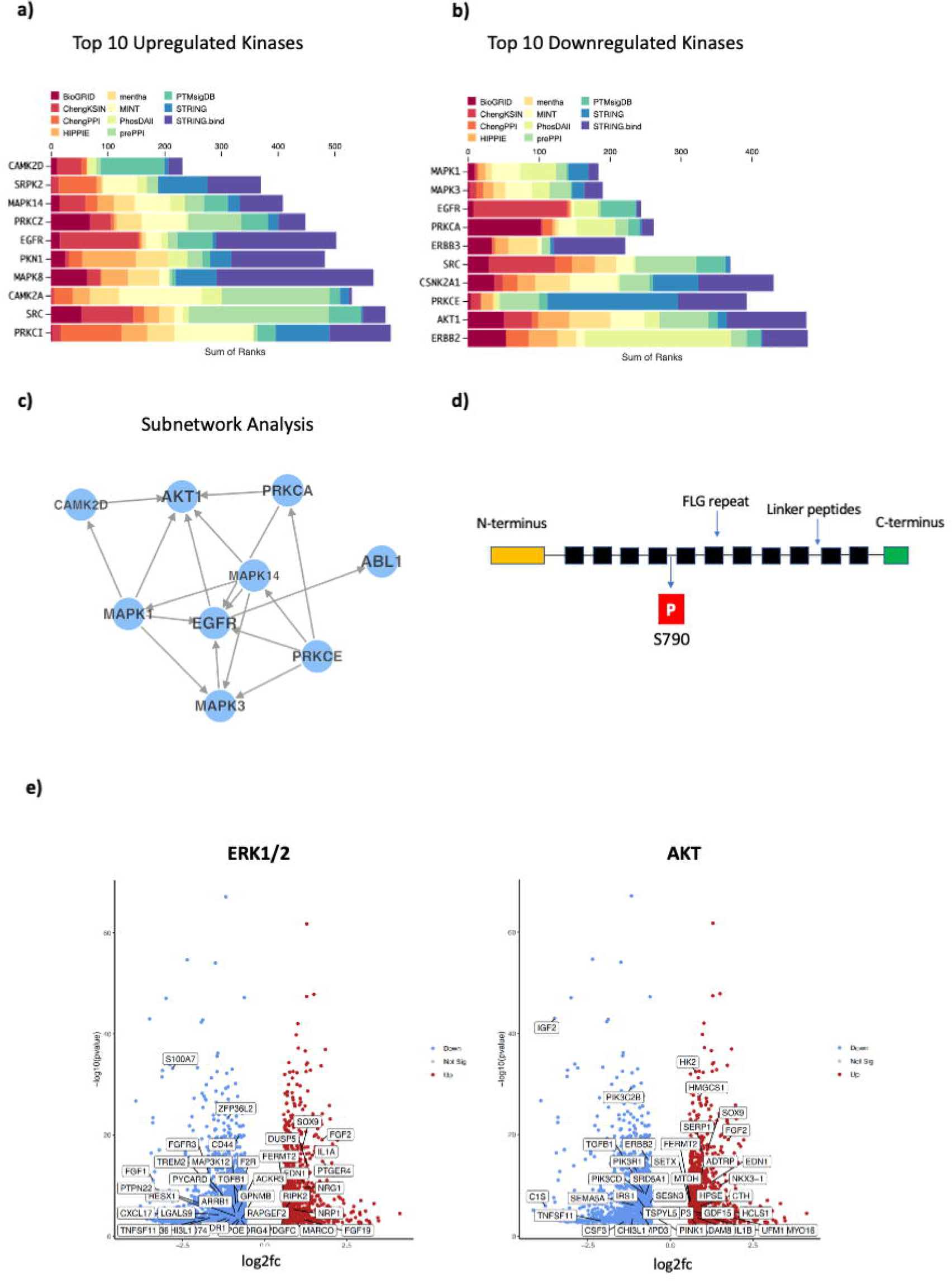
Phosphoproteomic Analysis Identifies Downregulation of the AKT and EKR1/2 Pathways and Upregulation of CAMK2D and the JNK Pathway Following FLG shRNA KD. a) Top 10 upregulated kinases using kinase enrichment analysis on phosphoroteomic data from FLG shRNA KD lysates. b) Top 10 Downregulated kinases. c) Subnetwork analysis of enriched kinases: directed edges indicate interactions supported by kinase-substrate evidence. d) Diagram showing the site where profilaggrin is differentially phosphorylated following FLG KD. e) Volcano plot showing differentially expressed ERK1/2 and AKT related genes using RNA-seq following FLG siRNA KD.

Kinase enrichment analysis (KEA) was used to predict upstream kinases and signalling pathways responsible for the differential phosphorylation state (Figure 4a-c). ERK1/2 were the most downregulated and CAMK2D the most upregulated kinase (Figure 4a-b). We were also interested in the downregulation of AKT1, given that AKT1 has an important role in terminal differentiation and FLG processing (Gutowska-Owsiak et al., 2018; Naeem et al., 2017; O’Shaughnessy et al., 2007). CAMK2D belongs to the Ca2+/calmodulin-dependent protein kinase II (CAMK2) family. Profilaggrin is a predicted target for CAMK2 using phosphorylation-site prediction software (Sandilands et al., 2009) but this has not been explored mechanistically. Chen et al. identified CAMK2D as a differentially expressed gene when comparing lesional and non-lesional AE skin to healthy controls using a pooled analysis of microarray datasets (L. H. Chen et al., 2022). However, CAMK2D is poorly enriched in the skin, so further work is necessary to establish if this protein has a role in barrier function or AE.

We did not see any enrichment of BMP or TGF-beta kinases from the phosphoproteomic dataset. However, the dataset did suggest mechanisms by which BMP-SMAD signalling is altered. Reduced ERK1/2 activation increases nuclear SMAD1/5/9 translocation, via phosphorylation of SMAD1/5/9 at a different phosphosite to the BMP receptor (Kretzschmar et al., 1997). Additionally, p38 and JNK, both of which are activated by BMP ligands via non-canonical BMP-signalling (Botchkarev & Sharov, 2004; Singh et al., 2012), were upregulated following FLG KD. This suggests that FLG KD leads to upregulation of non-canonical BMP signalling and that this pathway may be more relevant in keratinocytes. Kinase enrichment also favours the most studied kinases and underestimates pathways with a smaller number of substrates (Franciosa et al., 2023). BMP signalling may therefore have subsequently been underrepresented in the analysis.

Further analysis of our RNA-seq datasets confirmed dysregulation of ERK1/2 and AKT related genes following FLG KD (Figure 4e). Western blot analysis of FLG KD lysates confirmed downregulation of pAKT and the downstream protease Cathepsin H following FLG KD (Naeem et al., 2017; Figure 5a). Although ERK1/2 was not consistently downregulated (Figure 5a), AE patient skin biopsies showed a positive correlation between FLG and pAKT and pERK1/2 fluorescence (Figure 5b), confirming that this association existed *in vivo*. AKT1 is required for optimal FLG processing via Cathepsin H (Gutowska-Owsiak et al., 2018; Naeem et al., 2017; O’Shaughnessy et al., 2007). Our data suggests that loss of FLG caused a positive feedback loop with AKT1, whereby reduced AKT1 signalling lead to further reduction of FLG expression.

**Figure 5:**
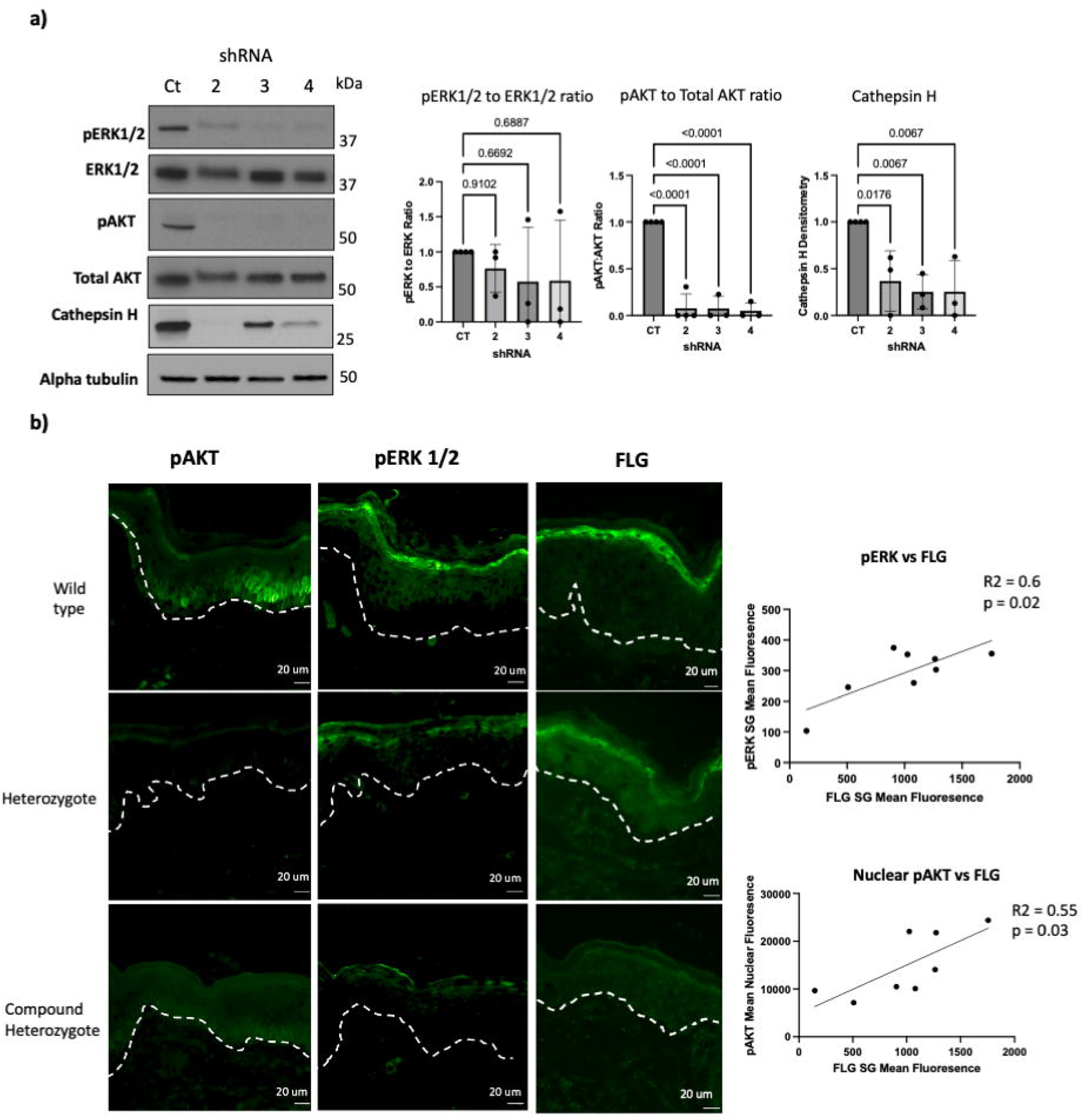
Loss of FLG is Associated with a Reduction in ERK1/2 and AKT1 Signalling. a) Western blots from FLG shRNA KD cell lines for pERK and pAKT and the AKT-downstream protease Cathepsin H b) Graphs of densitometry corresponding to the western blots. c) Staining for pERK1/2, pAKT1/2 and FLG using AE patient biopsies, with scatter plots demonstrating the relationship between phospho-kinase expression and FLG expression levels.

We next looked to establish a downstream effector that could link the BMP, AKT and ERK pathways. FOS was the fifth-most downregulated BMP related gene in the FLG RNA-seq KD dataset (Figure 2a). cFOS, the AP-1 transcription factor derived from the FOS gene, is induced by AKT1, ERK1/2 and JNK signalling (Wee & Wang, 2017; Wei & Liu, 2002). On review of our *in silico* analysis of BMP related genes, FOS positively correlated with FLG in populations where BMP6 or FST negatively correlated with FLG (Figure 2b). Further analysis of AP-1 related genes using the FLG KD RNA-seq dataset identified FOS and other AP-1 related genes as differentially expressed following FLG KD (Figure 6a). We confirmed a reduction in FOS expression following FLG KD using qPCR and western blotting on the FLG shRNA KD lysates (Figure 6b). Transcription factor enrichment analysis of the FLG KD RNA-seq dataset identified SMAD4 and FOSL2 as within the top 10 enriched transcription factors following FLG KD (Figure 6c). FOS positively correlated with FLG expression and was reduced in AE sections, (Figure 6d). rBMP6 treatment caused a reduction in cFOS expression (Figure 6e), but neither pAKT nor pERK1/2 expression were reduced (Supplementary Figure S4b). This suggests that while BMP signalling caused a reduction in cFOS expression, this was independent of both AKT and ERK1/2.

**Figure 6:**
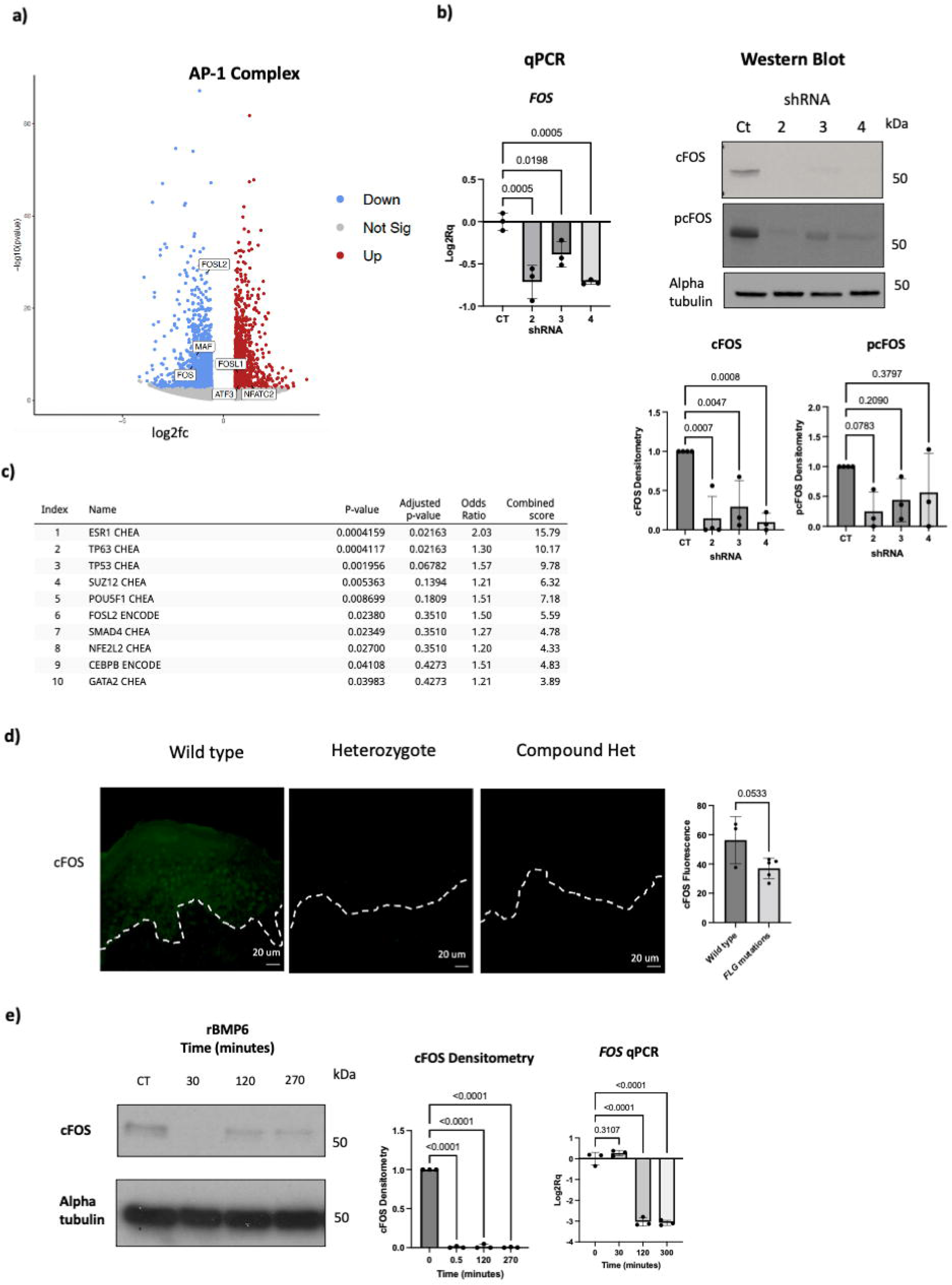
cFOS Expression is Reduced Following FLG Knock Down, which is Potentially Mediated by BMP6 Expression. a) Volcano plot for AP-1 related genes using RNA-seq following FLG siRNA KD in NHEKs. b) qPCR for FOS (expressed as Log2Rq relative to control. c) western blots for cFOS and p-cFOS following FLG shRNA KD in NHEKs. d) Transcription factor enrichment for the RNA-seq differentially expressed genes following FLG siRNA KD derived from the ENCODE and ChIP-X databases using Enrichr software. Index represents the rank, name the gene symbol and corresponding database it was derived from, p-value calculated by the Fisher exact test, adjusted p-value using the Benjamini-Hochberg method for multiple hypotheses testing, odds ratio represents deviation of the transcription factor from an expected rank of random gene sets, combined score is calculated as: log2p-value x the odds ratio. e) Tissue sections from AE patients with staining for cFOS. f) Western blot showing cFOS expression and qPCR showing FOS expression (expressed as Log2Rq relative to control) following rBMP6 treatment in REKs.

### Filaggrin loss and increased SMAD signalling alter desmosomal expression *in vitro* and ***in vivo***

Functional enrichment analysis of differentially regulated phosphoproteins yielded significant over-representation of desmosomal proteins. 13/25 desmosomal proteins were significantly differentially phosphorylated (Figure 7a), however there was only modest changes in gene expression in the Filaggrin kd keratinocytes (Figure 7b). We found an increased protein yield from compound heterozygote AE patients, suggestive of a potential defect in desmosomal function (Figure 7c). We were therefore interested to identify further downstream targets of pSMAD1/5/9 activation in the skin barrier, and their relationship to desmosomes. We performed proteomic analysis using lysates extracted from AE patients’ TS. These patients had undergone *FLG* genotyping and western blots were performed on the same TS samples to categorise their expression of pSMAD1/5/9. We identified a subset of pSMAD1/5/9 high patients that had significantly increased expression of Desmoplakin (DSP), Plakophilin 1 (PKP1), Desmoglein 3 (DSG3) and Desmocollin 3 (DSC3) (Figure 7c and d), suggesting that the BMP pathway may control desmosomal expression in AE. We also found reduced DSP expression in FLG heterozygote patients relative to controls (Figure 7e and f). This suggests that loss of FLG leads to changes to desmosomal expression and structure, which may partially be mediated by BMP signalling.

**Figure 7:**
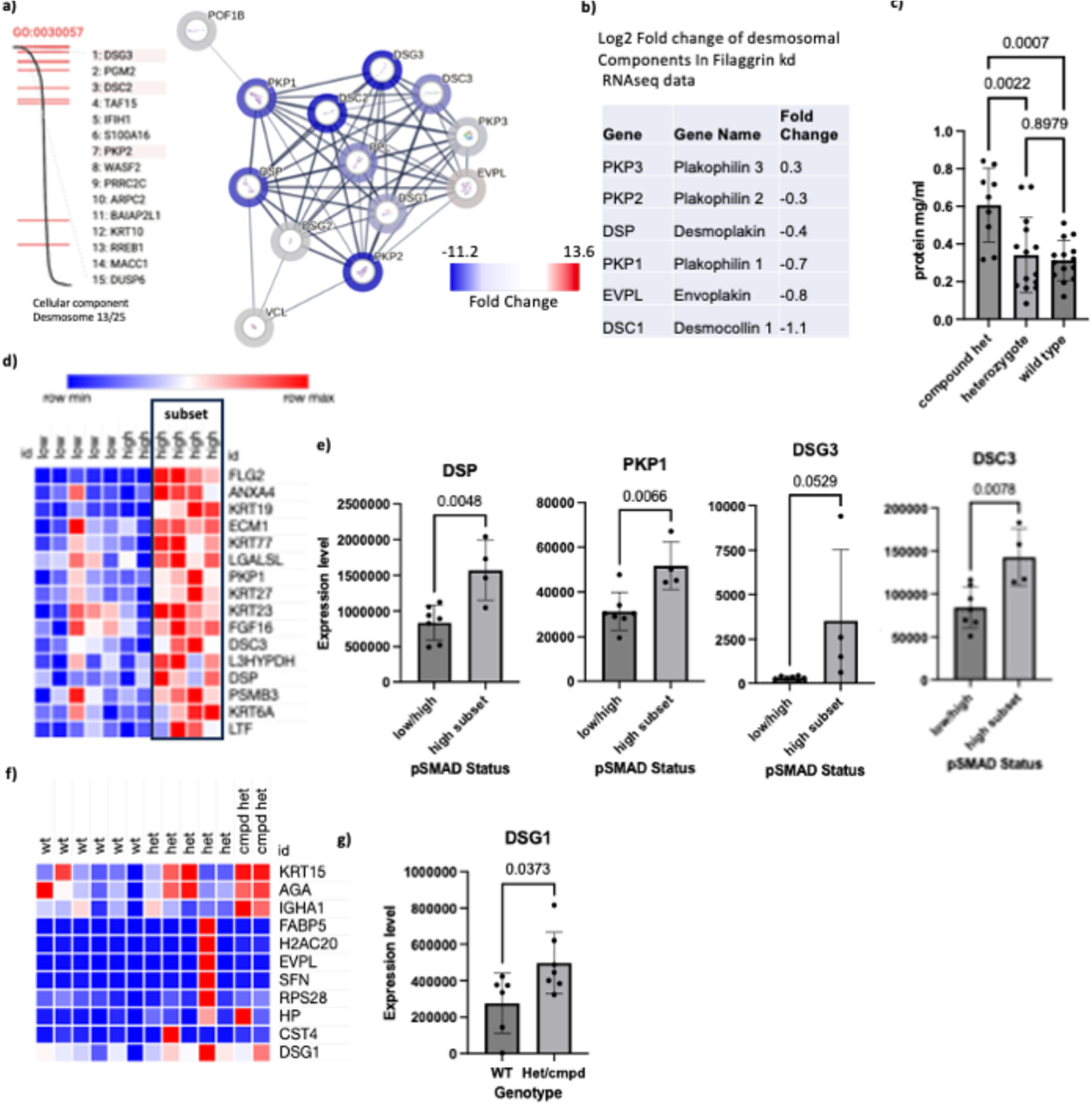
FLG Knock Down and increased SMAD1/5/9 signalling leads to Differential Phosphorylation and expression of Desmosomal Proteins. a) STRING analysis and functional enrichment analysis of desmosomal proteins in the phosphoproteomic analysis of FLG shRNA KD keratinocytes b) Differentially expressed desmosomal genes in the RNAseq analysis of FLG siRNA KD keratinocytes. c) Total protein yield from extracted AE patient tapestrips from different *FLG* genotypes. d) Heatmap of differentially expressed proteins from pSMAD 1/5/9 low and high AE patient tapestrips. The box indicates the subset of pSMAD 1/5/9 high patients with increased desmosomal expression. e) Graphs of expression of DSP, PKP1, DSG3 and DSC3 comparing the BMP high subset with the rest of the AE patients analysed. f) Heatmap of differentially expressed proteins from different FLG genotypes (het, heterozygote; cmpd het, compound heterozygote). g) Graphs of expression of DSG1 comparing heterozygotes and compound heterozygotes to wildtype.

## Discussion

Our data show that loss of FLG has a profound effect on the cellular environment with alteration of multiple signalling pathways. BMP signalling, with a subsequent reduction in cFOS expression was implicated as a key downstream pathway in patients with reduced FLG expression. Although BMP-TGF-beta kinases were not enriched using FLG KD phosphoproteomics, the increase in p38 and JNK pathways pathways suggests that the non-canonical BMP signalling pathway may be more relevant in keratinocytes. BMP6 activates the JNK pathway through phosphorylation but appears to reduce it through transcription. AKT1 and ERK1/2 both induce the transcription of cFOS (Wee & Wang, 2017), so reduced signalling in these pathways may be responsible for the changes to cFOS following FLG KD. Our data suggest that this is independent of BMP6, so further work is needed to tease out the ways that these pathways overlap in the keratinocyte cellular environment. ERK1/2 activation phosphorylates SMAD1 at a different site to the BMP-receptor (Kretzschmar et al., 1997). This has the opposite effect to BMP-triggered phosphorylation of SMAD1: SMAD1 is excluded from the nucleus, inhibiting its effects on gene expression (Kretzschmar et al., 1997). The reduction in ERK1/2 signalling seen following FLG KD may therefore be responsible for the increase in SMAD signalling.

There has been little research into the association between BMP signalling and AE. Naeem et al. showed an increase in nuclear SMAD1 expression in both lesional and non-lesional biopsies from AE patients (A. S. Naeem et al., 2015). Other groups demonstrated that activation of the TGF-beta pathway via SMAD2/3 signalling inhibits atopic inflammation (Anthoni et al., 2007; Kärner et al., 2017; Sumiyoshi et al., 2003). A genome wide association study in AE patients identified single nucleotide polymorphisms within SMAD4 and SMAD7 (Johansson et al., 2019): regulatory components of the BMP-TGF-beta signalling pathways.

Previous studies have shown that BMP2 negatively regulates FLG and EDC proteins in the epidermis (A. S. Naeem et al., 2015; Yu et al., 2010). Yu et al. found that constitutively active BMP receptor signalling or rBMP2 treatment of embryonic mice caused an ichthyotic phenotype with reduced FLG expression (Yu et al., 2010). The authors hypothesised that BMP negatively regulates FLG in the lower epidermis to prevent aberrant FLG expression and that a reduction of BMP in the stratum granulosum facilitates FLG expression at the correct level of the epidermis (Yu et al., 2010). Our data, and that of other authors, showed an increase in terminal differentiation proteins following rBMP6 treatment (Gosselet et al., 2007; Kim et al., 2018; McDonnell et al., 2001; Phillips et al., 2013). The positive correlation we saw between pSMAD1/5/9 and KRT2 in AE patients suggests that BMP signalling also increases suprabasal keratin expression *in vivo*. This suggests that BMP signalling has a protective role in response to barrier dysfunction. Strong and uniform BMP6 expression caused repression of keratinocyte proliferation, whereas moderate expression of BMP6 induced keratinocyte proliferation, suggesting that consistent with BMP’s morphogenic role during development, the effect of BMP signalling is both spatially, temporally and dose-dependent (Blessing et al., 1996;Botchkarev, 2003; Botchkarev & Sharov, 2004). After wounding there is BMP6 upregulation at the edge of the wound, followed by downregulation (Kaiser et al., 1998), whereas mice overexpressing BMP6 have delayed wound healing (Kaiser et al., 1998). This leads us to hypothesise that increased BMP signalling in AE may be a response to barrier injury.

No previous work has used phosphoproteomic analysis on FLG KD or AE samples. Two groups identified differential expression of the mTOR-AKT1 axis using proteomic analysis (Elias et al., 2017; Elias, Wright, Nicholson, et al., 2019), and reduction in pAKT and pERK1/2 following FLG KD (S. Wang et al., 2018). FLG co-localises with AKT1 in post confluent keratinocytes (Rogerson et al., 2021) and AKT is essential for FLG processing via the protease Cathepsin H (Gutowska-Owsiak et al., 2018; Aishath S. Naeem et al., 2017; O’Shaughnessy et al., 2007). In AE patients, *FLG* gene expression negatively correlates with *RPTOR*, which encodes RAPTOR, a component of the AKT signalling pathway (Aishath S. Naeem et al., 2017). Both the AKT1 and AKT2 knockout mice show reduced FLG expression and have a subtle cornified envelope defect (O’Shaughnessy et al., 2007). AKT1 phosphorylates EMSY, a transcriptional regulator of FLG, which is over-expressed in AE skin (Elias, Wright, Remenyi, et al., 2019). AKT1 phosphorylation of HspB1 allows FLG keratohyalin granules to dissociate from the actin scaffold, facilitating the release of processed FLG into the cytoplasm (Gutowska-Owsiak et al., 2018). Our analysis showed changes to the phosphorylation of the actin cytoskeleton regulators, suggesting that this AKT1-HspB1-FLG-Actin axis is disrupted following FLG KD. Taken together with our data, we propose a model in which FLG and AKT1 are in a positive feedback loop with each other (Figure 8). Loss of FLG leads to reduced AKT1 signalling, which further impairs FLG expression and AKT1 signalling. Finding ways to breaking this positive feedback loop may be a key mechanism to restore barrier function in AE.

**Figure 8:** Models Summarising Differential Phosphorylation of the FLG-AKT Axis Following FLG KD, and the effect of FLG KD on the cell signalling. a) 1. Inactive profilaggrin exists as keratohyalin granules. 2. These bind to the actin cytoskeleton where they translocate towards the nucleus. 3. AKT1, phosphorylates HspB1, enabling keratohyalin granules to dissociate from the actin scaffold. 4. Profilaggrin is metabolised to active FLG monomers via AKT1 and Cathepsin H. 5. AKT1 phosphorylates EMSY, a transcriptional regulator of FLG. 6. AKT1 phosphorylates nuclear lamins, leading to nuclear degradation and terminal differentiation (A. S. Naeem et al., 2015). AKT1, actin regulators and the profilaggrin protein are differentially phosphorylated following FLG KD. Our proposed model is that FLG and AKT1 are in a positive feedback loop with each other. Loss of FLG leads to reduced AKT1 signalling, which further impairs FLG expression and AKT1 signalling. b) Loss of FLG leads to increased BMP, JNK, p38 and CAMK2D signalling, leading to nuclear SMAD1 translocation, reduced cFOS expression and increased EDC/Keratin expression. Increased FST reduces TGF-Beta signalling and provides more sites for BMP6/2 to bind to the BMP receptor. Reduced AKT1 and ERK1/2 signalling, possibly mediated via the EGFr receptor, leads to reduced cFOS expression, and increased nuclear SMAD1 independent of BMP signalling. Reduced AKT1 expression leads to reduced FLG expression with a positive feedback loop.

FLG KD led to a global reduction in EDC and keratin gene expression. *In vivo*, FLG expression positively correlated with EDC and KRT gene expression across multiple datasets, but this was not replicated using protein analysis from our AE patient cohort. The disconnect between gene and protein expression warrants further study. The literature demonstrates conflicting results following FLG KD, with some studies showing a reduction (Pendaries et al., 2014; S. Wang et al., 2018; X. W. Wang et al., 2017) and others no change in terminal differentiation proteins (Elias, Wright, Nicholson, et al., 2019; Mildner et al., 2010). The reduction in keratin phosphorylation in FLG KD led us to hypothesise that a reduction in keratin expression following FLG KD also reflects a change to keratin phosphorylation. Phosphorylation of keratins is thought to affect keratin organisation, solubility and interactions (Loschke et al., 2015; Yang et al., 2012).

Components of the desmosome were highly enriched in our analysis. In the literature, electron microscopy of FLG KD organotypic models found ultrastructural changes to the desmosomes and corneodesmosomes (Elias, Wright, Nicholson, et al., 2019; Ipponjima et al., 2020), but no changes to desmosomal protein expression (Elias, Wright, Nicholson, et al., 2019). Ultrastructural changes without an overall change in desmosomal protein expression could be explained by the phosphorylation changes described herein. Given that FLG mutations predispose to food allergy (Brough et al., 2014) and the genetic disease SAM (severe dermatitis, multiple allergies and metabolic wasting) is due to mutations in desmoglein-1 (Bin & Leung, 2016), the desmosome offers a novel and interesting therapeutic target for AE. DSG3 is often upregulated in when DSG1 are lost, including the overactivity of serine proteases seen in the severe-AE-like genetic disease Netherton Syndrome, where loss of the LEKTI serine protease inhibitor leads to hyperactivity of Serine proteases, increased degradation of DSG1, and in response up-regulation of DSG3 (Zhu et al., 2017). BMP signalling may be inducing a similar response in AE.

Overall, our data show that loss of FLG has a profound effect on the cellular environment with changes to intracellular signalling pathways. We have identified key targets downstream of FLG, including AKT1, ERK1/2, cFOS, BMP6, the desmosome and CAMK2D, all of which could represent novel therapeutic targets to restore epidermal barrier function in AE.

## References

1. Abuabara, K., Ye, M., McCulloch, C. E., Sullivan, A., Margolis, D. J., Strachan, D. P., Paternoster, L., Yew, Y. W., Williams, H. C., & Langan, S. M. (2019). Clinical onset of atopic eczema: Results from 2 nationally representative British birth cohorts followed through midlife. Journal of Allergy and Clinical Immunology, 144(3), 710–719. 10.1016/j.jaci.2019.05.040

2. Alcolea, M. P., Casado, P., Rodríguez-Prados, J.-C., Vanhaesebroeck, B., & Cutillas, P. R. (2012). Phosphoproteomic analysis of leukemia cells under basal and drug-treated conditions identifies markers of kinase pathway activation and mechanisms of resistance. Molecular & Cellular Proteomics : MCP, 11(8), 453–466. 10.1074/mcp.M112.017483

3. Anthoni, M., Wang, G., Deng, C., Wolff, H. J., Lauerma, A. I., & Alenius, H. T. (2007). Smad3 signal transducer regulates skin inflammation and specific IgE response in murine model of atopic dermatitis. Journal of Investigative Dermatology, 127(8), 1923–1929. 10.1038/sj.jid.5700809

4. Bikle, D. D., Gee, E., & Pillai, S. (1993). Regulation of keratinocyte growth, differentiation, and vitamin D metabolism by analogs of 1,25-dihydroxyvitamin D. Journal of Investigative Dermatology, 101(5), 713–718. 10.1111/1523-1747.ep12371681

5. Bin, L., & Leung, D. Y. M. (2016). Genetic and epigenetic studies of atopic dermatitis. Allergy, Asthma and Clinical Immunology, 12(1), 1–14. 10.1186/s13223-016-0158-5

6. BioRender. (n.d.). BioRender. https://app.biorender.com/biorender-templates

7. Blessing, M., Schirmacher, P., & Kaiser, S. (1996). Overexpression of bone morphogenetic protein-6 (BMP-6) in the epidermis of transgenic mice: inhibition or stimulation of proliferation depending on the pattern of transgene expression and formation of psoriatic lesions. Journal of Cell Biology, 135(1), 227–239. 10.1083/jcb.135.1.227

8. Botchkarev, V. A. (2003). Bone morphogenetic proteins and their antagonists in skin and hair follicle biology. Journal of Investigative Dermatology, 120(1), 36–47. 10.1046/j.1523-1747.2003.12002.x

9. Botchkarev, V. A., & Sharov, A. A. (2004). BMP signaling in the control of skin development and hair follicle growth. Differentiation, 72(9–10), 512–526. 10.1111/j.1432-0436.2004.07209005.x

10. Brough, H. A., Simpson, A., Makinson, K., Hankinson, J., Brown, S., Douiri, A., Belgrave, D. C. M., Penagos, M., Stephens, A. C., McLean, W. H. I., Turcanu, V., Nicolaou, N., Custovic, A., & Lack, G. (2014). Peanut allergy: Effect of environmental peanut exposure in children with filaggrin loss-of-function mutations. Journal of Allergy and Clinical Immunology, 134(4), 867–875.e1. 10.1016/j.jaci.2014.08.011

11. Brown, S. J., Relton, C. L., Liao, H., Zhao, Y., Sandilands, A., McLean, W. H. I., Cordell, H. J., & Reynolds, N. J. (2009). Filaggrin haploinsufficiency is highly penetrant and is associated with increased severity of eczema: Further delineation of the skin phenotype in a prospective epidemiological study of 792 school children. British Journal of Dermatology, 161(4), 884–889. 10.1111/j.1365-2133.2009.09339.x

12. Cau, L., Pendaries, V., Lhuillier, E., Thompson, P. R., Serre, G., Takahara, H., Méchin, M. C., & Simon, M. (2017). Lowering relative humidity level increases epidermal protein deimination and drives human filaggrin breakdown. Journal of Dermatological Science, 86(2), 106–113. 10.1016/j.jdermsci.2017.02.280

13. Chen, E. Y., Tan, C. M., Kou, Y., Duan, Q., Wang, Z., Meirelles, G. V., Clark, N. R., & Ma’ayan, A. (2013). Enrichr: interactive and collaborative HTML5 gene list enrichment analysis tool. BMC Bioinformatics, 14(1), 128. 10.1186/1471-2105-14-128

14. Chen, L. H., Qi, X., Wang, J. Y., Yin, J. L., Sun, P. H., Sun, Y., Wu, Y., Zhang, L., & Gao, X. H. (2022). Identification of novel candidate genes and predicted miRNAs in atopic dermatitis patients by bioinformatic methods. Scientific Reports, 12(1), 1–10. 10.1038/s41598-022-26689-8

15. Choi, J. K., Yu, U., Kim, S., & Yoo, O. J. (2003). Combining multiple microarray studies and modeling interstudy variation. Bioinformatics, 19(SUPPL. 1). 10.1093/bioinformatics/btg1010

16. Czarnowicki, T., He, H., Krueger, J. G., & Guttman-Yassky, E. (2019). Atopic dermatitis endotypes and implications for targeted therapeutics. Journal of Allergy and Clinical Immunology, 143(1), 1–11. 10.1016/j.jaci.2018.10.032

17. Dale, B. A., Presland, R. B., Lewis, S. P., Underwood, R. A., & Fleckman, P. (1997). Transient expression of epidermal filaggrin in cultured cells causes collapse of intermediate filament networks with alteration of cell shape and nuclear integrity. Journal of Investigative Dermatology, 108(2), 179–187. 10.1111/1523-1747.ep12334205

18. Darling, A. L., Hart, K. H., Macdonald, H. M., Horton, K., Kang’ombe, A. R., Berry, J. L., & Lanham-New, S. A. (2013). Vitamin D deficiency in UK South Asian Women of childbearing age: a comparative longitudinal investigation with UK Caucasian women. Osteoporosis International : A Journal Established as Result of Cooperation between the European Foundation for Osteoporosis and the National Osteoporosis Foundation of the USA, 24(2), 477–488. 10.1007/s00198-012-1973-2

19. de Veer, S. J., Furio, L., Harris, J. M., & Hovnanian, A. (2014). Proteases: Common culprits in human skin disorders. Trends in Molecular Medicine, 20(3), 166–178. 10.1016/j.molmed.2013.11.005

20. Drislane, C., & Irvine, A. D. (2020). The role of filaggrin in atopic dermatitis and allergic disease. Annals of Allergy, Asthma and Immunology, 124(1), 36–43. 10.1016/j.anai.2019.10.008

21. Elias, M. S., Long, H. A., Newman, C. F., Wilson, P. A., West, A., McGill, P. J., Wu, K. C., Donaldson, M. J., & Reynolds, N. J. (2017). Proteomic analysis of filaggrin deficiency identifies molecular signatures characteristic of atopic eczema. Journal of Allergy and Clinical Immunology, 140(5), 1299–1309. 10.1016/j.jaci.2017.01.039

22. Elias, M. S., Wright, S. C., Nicholson, W. V., Morrison, K. D., Prescott, A. R., Have, S. Ten, Whitfield, P. D., Lamond, A. I., & Brown, S. J. (2019). Proteomic analysis of a filaggrin-deficient skin organoid model shows evidence of increased transcriptional-translational activity, keratinocyte-immune crosstalk and disordered axon guidance. Wellcome Open Research, 4, 1–21. 10.12688/wellcomeopenres.15405.1

23. Elias, M. S., Wright, S. C., Remenyi, J., Abbott, J. C., Bray, S. E., Cole, C., Edwards, S., Gierlinski, M., Glok, M., McGrath, J. A., Nicholson, W. V., Paternoster, L., Prescott, A. R., Have, S. Ten, Whitfield, P. D., Lamond, A. I., & Brown, S. J. (2019). EMSY expression affects multiple components of the skin barrier with relevance to atopic dermatitis. Journal of Allergy and Clinical Immunology, 144(2), 470–481. 10.1016/j.jaci.2019.05.024

24. Elmose, C., & Thomsen, S. F. (2015). Twin Studies of Atopic Dermatitis: Interpretations and Applications in the Filaggrin Era. Journal of Allergy, 2015, 1–7. 10.1155/2015/902359

25. Franciosa, G., Locard-Paulet, M., Jensen, L. J., & Olsen, J. V. (2023). Recent advances in kinase signaling network profiling by mass spectrometry. Current Opinion in Chemical Biology, 73, 102260. 10.1016/j.cbpa.2022.102260

26. Gosselet, F. P., Magnaldo, T., Culerrier, R. M., Sarasin, A., & Ehrhart, J. C. (2007). BMP2 and BMP6 control p57Kip2 expression and cell growth arrest/terminal differentiation in normal primary human epidermal keratinocytes. Cellular Signalling, 19(4), 731–739. 10.1016/j.cellsig.2006.09.006

27. Gutowska-Owsiak, D., De La Serna, J. B., Fritzsche, M., Naeem, A., Podobas, E. I., Leeming, M., Colin-York, H., O’Shaughnessy, R., Eggeling, C., & Ogg, G. S. (2018). Orchestrated control of filaggrin-actin scaffolds underpins cornification. Cell Death and Disease, 9(4). 10.1038/s41419-018-0407-2

28. Hällqvist, J., Pinto, R.C., Heywood, W.E., Cordey, J., Foulkes, A.J.M., Slattery, C.F., Leckey, C.A., Murphy, E.C., Zetterberg, H., Schott, J.M., Mills, K., Paterson, R.W.. A Multiplexed Urinary Biomarker Panel Has Potential for Alzheimer’s Disease Diagnosis Using Targeted Proteomics and Machine Learning. Int J Mol Sci. 2023 Sep 6;24(18):13758. doi: 10.3390/ijms241813758.

29. Henderson, J., Northstone, K., Lee, S. P., Liao, H., Zhao, Y., Pembrey, M., Mukhopadhyay, S., Smith, G. D., Palmer, C. N. A., McLean, W. H. I., & Irvine, A. D. (2008). The burden of disease associated with filaggrin mutations: A population-based, longitudinal birth cohort study. Journal of Allergy and Clinical Immunology, 121(4). 10.1016/j.jaci.2008.01.026

30. Hönzke, S., Wallmeyer, L., Ostrowski, A., Radbruch, M., Mundhenk, L., Schäfer-Korting, M., & Hedtrich, S. (2016). Influence of Th2 Cytokines on the Cornified Envelope, Tight Junction Proteins, and β-Defensins in Filaggrin-Deficient Skin Equivalents. Journal of Investigative Dermatology, 136(3), 631–639. 10.1016/j.jid.2015.11.007

31. Howell, M. D., Kim, B. E., Gao, P., Grant, A. V., Boguniewicz, M., DeBenedetto, A., Schneider, L., Beck, L. A., Barnes, K. C., & Leung, D. Y. M. (2009). Cytokine modulation of atopic dermatitis filaggrin skin expression. Journal of Allergy and Clinical Immunology, 124(3 SUPPL. 2), 7–12. 10.1016/j.jaci.2009.07.012

32. Ipponjima, S., Umino, Y., Nagayama, M., & Denda, M. (2020). Live imaging of alterations in cellular morphology and organelles during cornification using an epidermal equivalent model. Scientific Reports, 10(1), 1–12. 10.1038/s41598-020-62240-3

33. Irvine, A. D. (2014). Crossing barriers; restoring barriers? Filaggrin protein replacement takes a bow. Journal of Investigative Dermatology, 134(2), 313–314. 10.1038/jid.2013.506

34. Johansson, Å., Rask-Andersen, M., Karlsson, T., & Ek, W. E. (2019). Genome-wide association analysis of 350 000 Caucasians from the UK Biobank identifies novel loci for asthma, hay fever and eczema. Human Molecular Genetics, 28(23), 4022–4041. 10.1093/hmg/ddz175

35. Ju, J.-H., Yang, W., Lee, K.-M., Oh, S., Nam, K., Shim, S., Shin, S. Y., Gye, M. C., Chu, I.-S., & Shin, I. (2013). Regulation of cell proliferation and migration by keratin19-induced nuclear import of early growth response-1 in breast cancer cells. Clinical Cancer Research : An Official Journal of the American Association for Cancer Research, 19(16), 4335–4346. 10.1158/1078-0432.CCR-12-3295

36. Kaiser, S., Schirmacher, P., Philipp, A., Protschka, M., Moll, I., Nicol, K., & Blessing, M. (1998). Induction of bone morphogenetic protein-6 in skin wounds. Delayed reepitheliazation and scar formation in BMP-6 overexpressing transgenic mice. Journal of Investigative Dermatology, 111(6), 1145–1152. 10.1046/j.1523-1747.1998.00407.x

37. Kärner, J., Wawrzyniak, M., Tankov, S., Runnel, T., Aints, A., Kisand, K., Altraja, A., Kingo, K., Akdis, C. A., Akdis, M., & Rebane, A. (2017). Increased microRNA-323-3p in IL-22/IL-17-producing T cells and asthma: a role in the regulation of the TGF-β pathway and IL-22 production. Allergy: European Journal of Allergy and Clinical Immunology, 72(1), 55–65. 10.1111/all.12907

38. Kim, J. H., Bae, H. C., Kim, J., Lee, H., Ryu, W. I., Son, E. D., Lee, T. R., Jeong, S. H., & Son, S. W. (2018). HIF-1α-mediated BMP6 down-regulation leads to hyperproliferation and abnormal differentiation of keratinocytes in vitro. Experimental Dermatology, 27(11), 1287–1293. 10.1111/exd.13785

39. Kretzschmar, M., Doody, J., & Massagué, J. (1997). Opposing BMP and EGF signalling pathways converge on the TGF-β family mediator Smad1. Nature, 389(6651), 618–622. 10.1038/39348

40. Kuleshov, M. V., Xie, Z., London, A. B. K., Yang, J., Evangelista, J. E., Lachmann, A., Shu, I., Torre, D., & Ma’Ayan, A. (2021). KEA3: Improved kinase enrichment analysis via data integration. Nucleic Acids Research, 49(W1), W304–W316. 10.1093/nar/gkab359

41. Kypriotou, M., Huber, M., & Hohl, D. (2012). The human epidermal differentiation complex: Cornified envelope precursors, S100 proteins and the “fused genes” family. Experimental Dermatology, 21(9), 643–649. 10.1111/j.1600-0625.2012.01472.x

42. Loschke, F., Seltmann, K., Bouameur, J. E., & Magin, T. M. (2015). Regulation of keratin network organization. Current Opinion in Cell Biology, 32, 56–64. 10.1016/j.ceb.2014.12.006

43. Margolis, D. J., Mitra, N., Wubbenhorst, B., & Nathanson, K. L. (2020). Filaggrin sequencing and bioinformatics tools. Archives of Dermatological Research, 312(2), 155–158. 10.1007/s00403-019-01956-3

44. McDonnell, M. A., Law, B. K., Serra, R., & Moses, H. L. (2001). Antagonistic effects of TGF/β1 and BMP-6 on skin keratinocyte differentiation. Experimental Cell Research, 263(2), 265–273. 10.1006/excr.2000.5117

45. Mildner, M., Jin, J., Eckhart, L., Kezic, S., Gruber, F., Barresi, C., Stremnitzer, C., Buchberger, M., Mlitz, V., Ballaun, C., Sterniczky, B., Födinger, D., & Tschachler, E. (2010). Knockdown of filaggrin impairs diffusion barrier function and increases UV sensitivity in a human skin model. Journal of Investigative Dermatology, 130(9), 2286–2294. 10.1038/jid.2010.115

46. Montoya, A., Beltran, L., Casado, P., Rodríguez-Prados, J.-C., & Cutillas, P. R. (2011). Characterization of a TiO₂ enrichment method for label-free quantitative phosphoproteomics. Methods (San Diego, Calif.), 54(4), 370–378. 10.1016/j.ymeth.2011.02.004

47. Morikawa, M., Koinuma, D., Tsutsumi, S., Vasilaki, E., Kanki, Y., Heldin, C. H., Aburatani, H., & Miyazono, K. (2011). ChIP-seq reveals cell type-specific binding patterns of BMP-specific Smads and a novel binding motif. Nucleic Acids Research, 39(20), 8712–8727. 10.1093/nar/gkr572

48. Moura, J., Da Silva, L., Cruz, M. T., & Carvalho, E. (2013). Molecular and cellular mechanisms of bone morphogenetic proteins and activins in the skin: Potential benefits for wound healing. Archives of Dermatological Research, 305(7), 557–569. 10.1007/s00403-013-1381-2

49. Naeem, A. S., Zhu, Y., Di, W. L., Marmiroli, S., & O’Shaughnessy, R. F. L. (2015). AKT1-mediated Lamin A/C degradation is required for nuclear degradation and normal epidermal terminal differentiation. Cell Death and Differentiation, 22(12), 2123–2132. 10.1038/cdd.2015.62

50. Naeem, Aishath S., Tommasi, C., Cole, C., Brown, S. J., Zhu, Y., Way, B., Willis Owen, S. A. G., Moffatt, M., Cookson, W. O., Harper, J. I., Di, W. L., Brown, S. J., Reinheckel, T., & O’Shaughnessy, R. F. L. (2017). A mechanistic target of rapamycin complex 1/2 (mTORC1)/V-Akt murine thymoma viral oncogene homolog 1 (AKT1)/cathepsin H axis controls filaggrin expression and processing in skin, a novel mechanism for skin barrier disruption in patients with atopic dermat. Journal of Allergy and Clinical Immunology, 139(4), 1228–1241. 10.1016/j.jaci.2016.09.052

51. O’Shaughnessy, R. F. L., Welti, J. C., Cooke, J. C., Avilion, A. A., Monks, B., Birnbaum, M. J., & Byrne, C. (2007). AKT-dependent HspB1 (Hsp27) activity in epidermal differentiation. Journal of Biological Chemistry, 282(23), 17297–17305. 10.1074/jbc.M610386200

52. Odhiambo, J. A., Williams, H. C., Clayton, T. O., Robertson, C. F., Asher, M. I., AÃ̅t-Khaled, N., Anderson, H. R., Beasley, R., Bj Ã– RkstÃ©n, B., Brunekreef, B., Crane, J., Ellwood, P., Flohr, C., Foliaki, S., Forastiere, F., GarcÃ¬a-Marcos, L., Keil, U., Lai, C. K. W., Mallol, J., … Nilsson, L. (2009). Global variations in prevalence of eczema symptoms in children from ISAAC Phase Three. Journal of Allergy and Clinical Immunology, 124(6), 1251–1258.e23. 10.1016/j.jaci.2009.10.009

53. Palmer, C. N. A., Irvine, A. D., Terron-Kwiatkowski, A., Zhao, Y., Liao, H., Lee, S. P., Goudie, D. R., Sandilands, A., Campbell, L. E., Smith, F. J. D., O’Regan, G. M., Watson, R. M., Cecil, J. E., Bale, S. J., Compton, J. G., DiGiovanna, J. J., Fleckman, P., Lewis-Jones, S., Arseculeratne, G., … McLean, W. H. I. (2006). Common loss-of-function variants of the epidermal barrier protein filaggrin are a major predisposing factor for atopic dermatitis. Nature Genetics, 38(4), 441–446. 10.1038/ng1767

54. Paternoster, L., Savenije, O. E. M., Heron, J., Evans, D. M., Vonk, J. M., Brunekreef, B., Wijga, A. H., Henderson, A. J., Koppelman, G. H., & Brown, S. J. (2018). Identification of atopic dermatitis subgroups in children from 2 longitudinal birth cohorts. Journal of Allergy and Clinical Immunology, 141(3), 964–971. 10.1016/j.jaci.2017.09.044

55. Pearton, D. J., Dale, B. A., & Presland, R. B. (2002). Functional analysis of the profilaggrin N-terminal peptide: Identification of domains that regulate nuclear and cytoplasmic distribution. Journal of Investigative Dermatology, 119(3), 661–669. 10.1046/j.1523-1747.2002.01831.x

56. Pendaries, V., Malaisse, J., Pellerin, L., Le Lamer, M., Nachat, R., Kezic, S., Schmitt, A. M., Paul, C., Poumay, Y., Serre, G., & Simon, M. (2014). Knockdown of filaggrin in a three-dimensional reconstructed human epidermis impairs keratinocyte differentiation. Journal of Investigative Dermatology, 134(12), 2938–2946. 10.1038/jid.2014.259

57. Phillips, M. A., Qin, Q., Hu, Q., Zhao, B., & Rice, R. H. (2013). Arsenite suppression of BMP signaling in human keratinocytes. Toxicology and Applied Pharmacology, 269(3), 290–296. 10.1016/j.taap.2013.02.017

58. Pigors, M., Common, J. E. A., Wong, X. F. C. C., Malik, S., Scott, C. A., Tabarra, N., Liany, H., Liu, J., Limviphuvadh, V., Maurer-Stroh, S., Tang, M. B. Y., Lench, N., Margolis, D. J., van Heel, D. A., Mein, C. A., Novak, N., Baurecht, H., Weidinger, S., McLean, W. H. I., … Kelsell, D. P. (2018). Exome Sequencing and Rare Variant Analysis Reveals Multiple Filaggrin Mutations in Bangladeshi Families with Atopic Eczema and Additional Risk Genes. Journal of Investigative Dermatology, 138(12), 2674–2677. 10.1016/j.jid.2018.05.013

59. Rieko, K.-K., & Motonobu, N. (2016). Effect of cis-urocanic acid on atopic dermatitis in NC/Nga mice. Journal of Dermatological Science, 84(1), e65–e66. 10.1016/j.jdermsci.2016.08.203

60. Rogerson, C., Wotherspoon, D. J., Tommasi, C., Button, R. W., & O’Shaughnessy, R. F. L. (2021). Akt1-associated actomyosin remodelling is required for nuclear lamina dispersal and nuclear shrinkage in epidermal terminal differentiation. Cell Death and Differentiation, 28(6), 1849–1864. 10.1038/s41418-020-00712-9

61. Sandilands, A., Sutherland, C., Irvine, A. D., & McLean, W. H. I. (2009). Filaggrin in the frontline: Role in skin barrier function and disease. Journal of Cell Science, 122(9), 1285– 1294. 10.1242/jcs.033969

62. Scott, I. R., & Harding, C. R. (1986). Filaggrin breakdown to water binding compounds during development of the rat stratum corneum is controlled by the water activity of the environment. Developmental Biology, 115(1), 84–92. 10.1016/0012-1606(86)90230-7

63. Simonsen, S., Thyssen, J. P., Heegaard, S., Kezic, S., & Skov, L. (2017). Expression of filaggrin and its degradation products in human skin following erythemal doses of ultraviolet B irradiation. Acta Dermato-Venereologica. 10.2340/00015555-2662

64. Singh, S. K., Abbas, W. A., & Tobin, D. J. (2012). Bone morphogenetic proteins differentially regulate pigmentation in human skin cells. Journal of Cell Science, 125(18), 4306–4319. 10.1242/jcs.102038

65. Smith, N., Sievert, L. L., Muttukrishna, S., Begum, K., Murphy, L., Sharmeen, T., Gunu, R., Chowdhury, O., & Bentley, G. R. (2021). Mismatch: a comparative study of vitamin D status in British-Bangladeshi migrants. Evolution, Medicine, and Public Health, 9(1), 164–173. 10.1093/emph/eoab001

66. Sumiyoshi, K., Nakao, A., Setoguchi, Y., Tsuboi, R., Okumura, K., & Ogawa, H. (2003). TGF-β/Smad signaling inhibits IFN-γ and TNF-α-induced TARC (CCL17) production in HaCaT cells. Journal of Dermatological Science, 31(1), 53–58. 10.1016/S0923-1811(02)00141-X

67. Tagoe, H., Hassan, S., Youssef, G., Heywood, W., Mills, K., Harper, J. I., & O’Shaughnessy, R. F. L. (2023). Chronic activation of Toll-like receptor 2 induces an ichthyotic skin phenotype. The British Journal of Dermatology. 10.1093/bjd/ljad095

68. Thomas, B. R., Tan, X. L., Van Duijvenboden, S., Hogan, S. C., Hughes, A. J., Tawfik, S. S., Dhoat, S., Atkar, R., Robinson, E. J., Rahman, S. R., Rahman, S., Ahmed, R. A., Begum, R., Khanam, H., Bourne, E. L., Wozniak, E. L., Mein, C. A., Kelsell, D. P., & O’Toole, E. A. (2023). Deep palmar phenotyping in atopic eczema: patterns associated with Filaggrin variants, disease severity and barrier function in a South Asian population. British Journal of Dermatology, 1–23. 10.1093/bjd/ljad036

69. Toomey, C.E., Heywood, W.E., Evans, J.R., Lachica, J., Pressey, S.N., Foti SC, Al Shahrani, M., D’Sa, K., Hargreaves, I.P., Heales, S., Orford, M., Troakes, C., Attems, J., Gelpi, E., Palkovits, M., Lashley, T., Gentleman, S.M., Revesz, T., Mills, K., Gandhi, S. Mitochondrial dysfunction is a key pathological driver of early stage Parkinson’s. Acta Neuropathol Commun. 2022 Sep 8;10(1):134. doi: 10.1186/s40478-022-01424-6.

70. Wang, S., Qiu, L., Meng, X., & Dang, N. (2018). Knock-down of filaggrin influences the mitogen-activated protein kinases signaling pathway in normal human epidermal keratinocytes. Medecine/Sciences, 34, 94–98. 10.1051/medsci/201834f116

71. Wang, X. W., Wang, J. J., Gutowska-Owsiak, D., Salimi, M., Selvakumar, T. A., Gwela, A., Chen, L. Y., Wang, Y. J., Giannoulatou, E., & Ogg, G. (2017). Deficiency of filaggrin regulates endogenous cysteine protease activity, leading to impaired skin barrier function. Clinical and Experimental Dermatology, 42(6), 622–631. 10.1111/ced.13113

72. Wee, P., & Wang, Z. (2017). Epidermal growth factor receptor cell proliferation signaling pathways. Cancers, 9(5), 1–45. 10.3390/cancers9050052

73. Wei, Z., & Liu, H. T. (2002). MAPK signal pathways in the regulation of cell proliferation in mammalian cells. Cell Research, 12(1), 9–18. 10.1038/sj.cr.7290105

74. Yang, F., Waters, K. M., Webb-Robertson, B.-J., Sowa, M. B., von Neubeck, C., Aldrich, J. T., Markillie, L. M., Wirgau, R. M., Gritsenko, M. A., Zhao, R., Camp, D. G., Smith, R. D., & Stenoien, D. L. (2012). Quantitative phosphoproteomics identifies filaggrin and other targets of ionizing radiation in a human skin model. Experimental Dermatology, 21(5), 352–357. 10.1111/j.1600-0625.2012.01470.x

75. Yu, X., Espinoza-Lewis, R. A., Sun, C., Lin, L., He, F., Xiong, W., Yang, J., Wang, A., & Chen, Y. (2010). Overexpression of constitutively active BMP-receptor-IB in mouse skin causes an ichthyosis-vulgaris-like disease. Cell and Tissue Research, 342(3), 401–410. 10.1007/s00441-010-1077-2

76. Zhu, Y., Underwood, J., Macmillan, D., Shariff, L., O’Shaughnessy, R., Harper, J.I., Pickard, C., Friedmann, P.S., Healy, E., Di, W.L., (2017) Persistent kallikrein 5 activation induces atopic dermatitis-like skin architecture independent of PAR2 activity. J Allergy Clin Immunol. 140(5):1310–1322

