## Supplementary Information for "Reduced Filaggrin expression induces dysregulated intracellular signalling in atopic eczema"

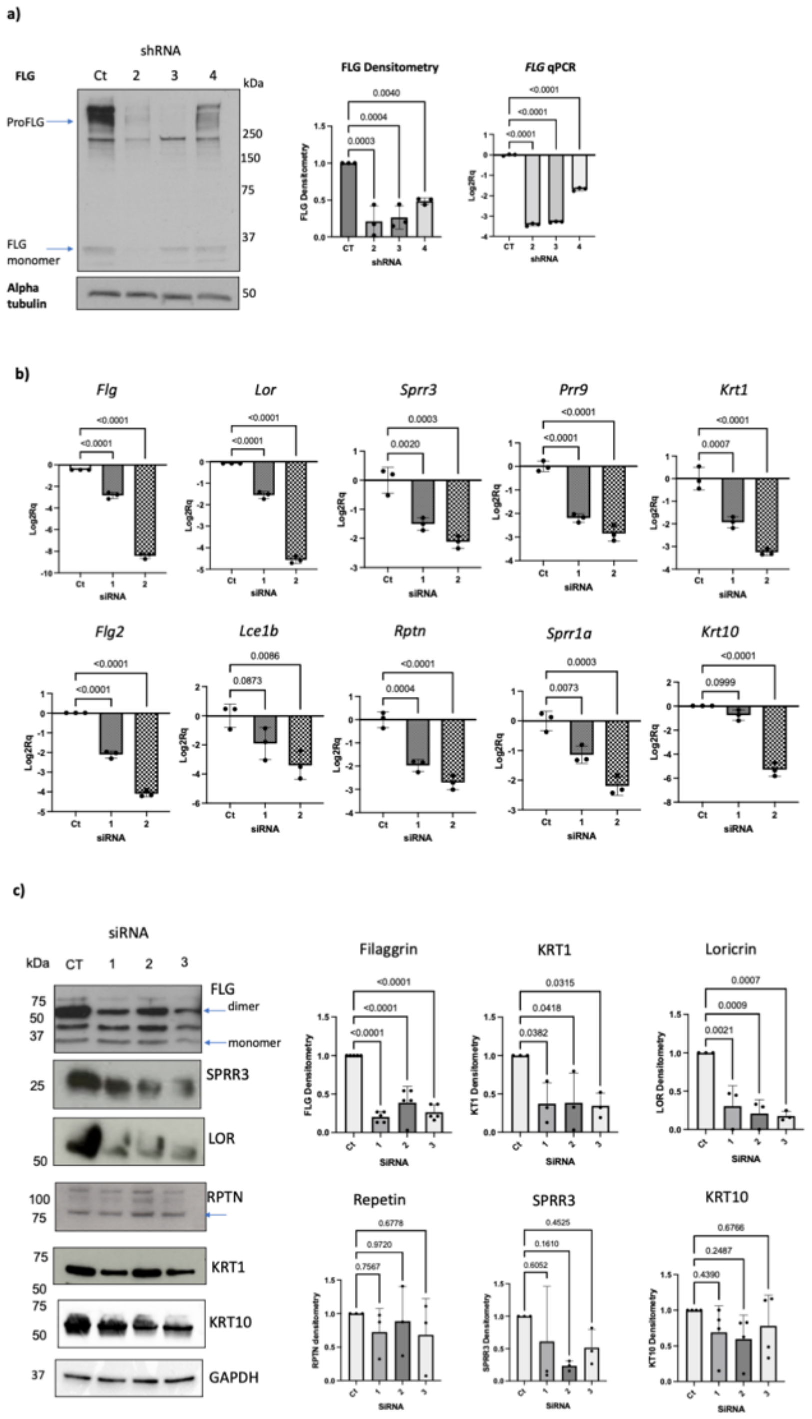


**Supplementary Figure S1:**

a) Western blots and qPCR following FLG shRNA KD in NHEKs confirming KD of FLG following shRNA transfection. b) qPCR following Flg siRNA KD in REKs expressed as Log2Rq relative to control. c) Western blots following Flg siRNA KD in NHEKs (representative of 3 biological replicates).

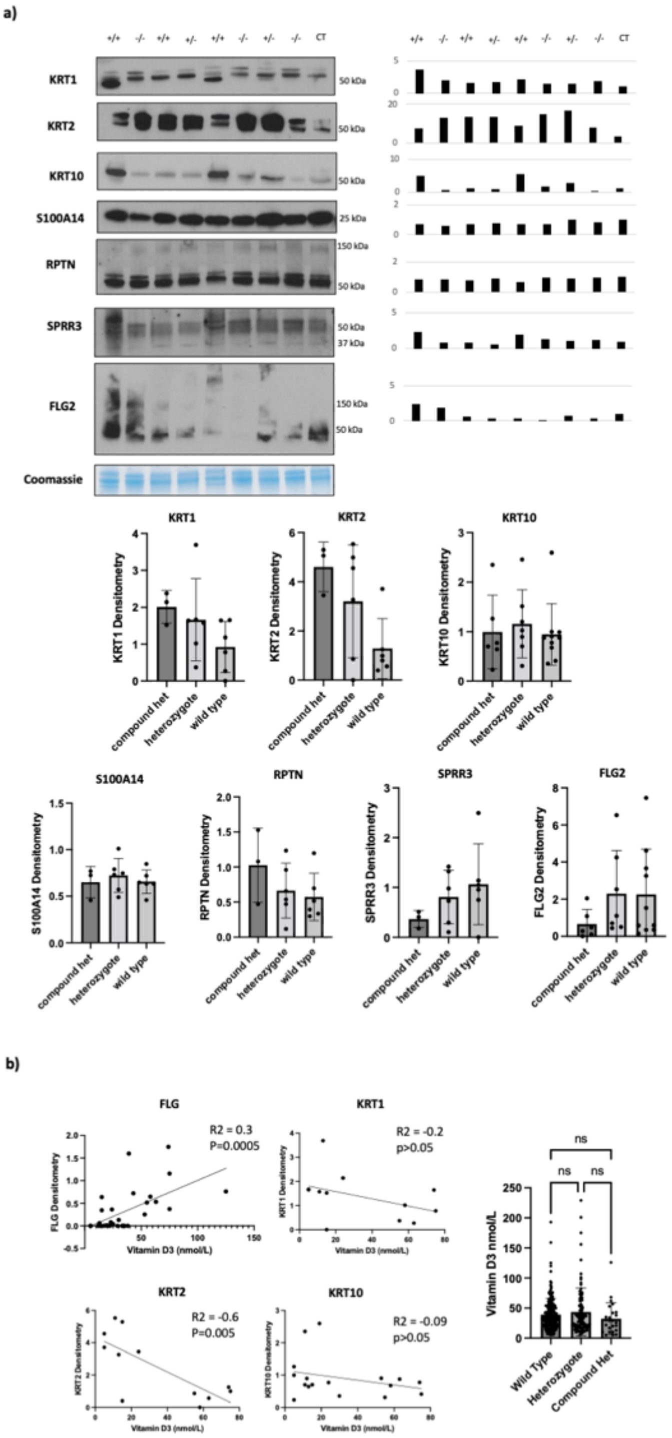


**Supplementary Figure S2:**

a) Western blots using AE patient tape strip lysates for EDC and suprabasal proteins. +/+ = wild type, +/- = heterozygote, -/- = compound heterozygote, with bar charts showing no association by genotype. b) Scatter plots demonstrating correlations between serum vitamin D3 (nearest level) and FLG or suprabasal keratin expression (from TS western blot data). Bar chart demonstrating no difference in serum vitamin D3 level and FLG genotype.

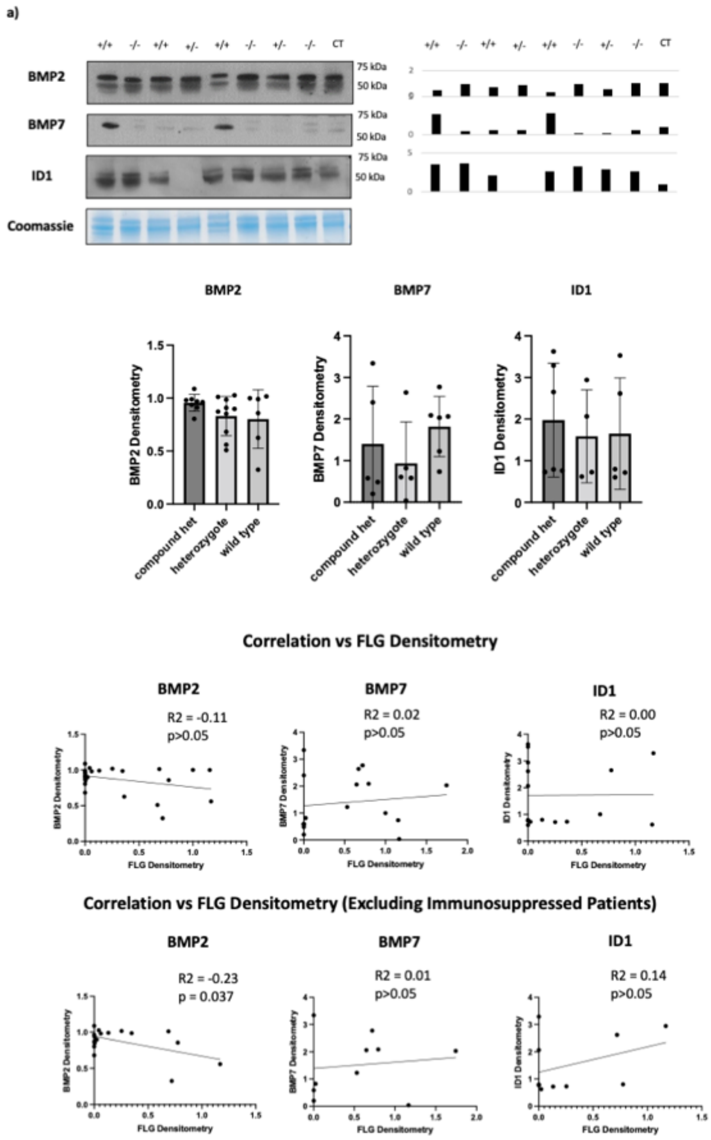


**Supplementary Figure S3:**

a) Western blots using AE patient tape strip lysates for BMP2, BMP7 and ID1. +/+ = wild type, +/- = heterozygote, -/- = compound heterozygote. Bar charts showing no association by genotype. Scatter plots demonstrating no correlation with FLG densitometry in all patients, but a negative correlation between FLG and BMP2 when immunosuppressed patients were excluded.

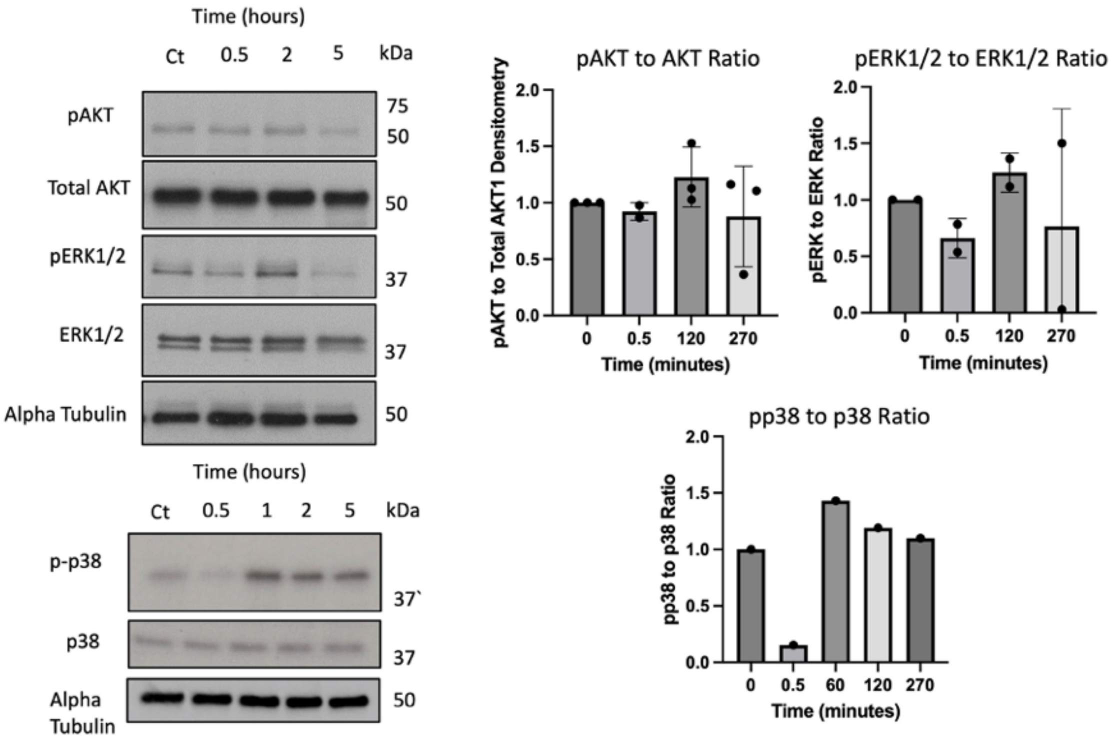


**Supplementary Figure S4:**

a) STRING analysis of the FLG shRNA KD Phosphoproteomic Dataset. Colours correspond to gene ontology analysis: yellow = keratinocyte differentiation, turquoise = desmosomes, green = regulation of actin filament-based process, blue = protein serine/threonine kinase activity, pink = microtubule cytoskeleton, red = keratin filament. b) Western blots showing no change in pAKT or PERK1/2 and an increase in pp38 expression following treatment of REKs with recombinant BMP6.

**Supplementary Table ST1:**

DSeq analysis of FLG siRNA knockdown of significant RNAseq data. baseMean, average of the normalized count values. Log2Fold Change, log transformed average fold changed of FLG siRNA compared to controls (n=3 each). Padj, FDR adjusted p-Value of the comparison. geneName, transcript ID.

**Supplementary Table ST2:**

File of all differentially expressed phosphoproteins in two different FLG shRNA knockdowns. Acc_no, Uniprot Accession number. Av FC, average fold change over both FLG shRNA knockdown lines. Av pval, average corrected p-value (FDR) across both lines.

**Supplementary Table ST3:**

Progenesis-derived Expression data for proteomic analysis of Eczema tapestrip samples sorted on the basis of pSMAD expression (pSMAD median), Filaggrin levels (Filag) or *FLG* genotype (wt, wildtype, het, heterozygous, cmpd het, compound heterozygous). Peptide count, number of peptides identified, unique peptides, number of ungrouped peptides for that protein. Confidence score is confidence in the protein assignment based on number of grouped and unique peptides.
